# Field size as a predictor of “excellence.” The selection of subject fields in Germany’s Excellence Initiative

**DOI:** 10.1101/2024.03.06.583816

**Authors:** Thomas Heinze, Isabel M. Habicht, Paul Eberhardt, Dirk Tunger

**Affiliations:** Institute of Sociology, University of Wuppertal, Wuppertal, Germany; Project Management, Research Center Jülich, Germany

**Keywords:** Research Excellence, Subject Field, Research Funding, Matthew Effect, Quantitative Analysis

## Abstract

We investigate the selection of subject fields in Germany’s “excellence initiative,” a two-phase funding scheme administered by the German Research Foundation (DFG) from 2005 to 2017 to increase international competitiveness of scientific research at German universities. While most empirical studies have examined the “excellence initiative’s” effects at the university level (“elite universities”), we focus on subject fields within universities. Based on both descriptive and logistic regression analyses, we find that the “excellence initiative” reveals a stable social order of public universities based on organizational size, that field selection is biased toward those fields with many professors and considerable grant funding, and that funding success in the second phase largely follows decisions from the first phase. We discuss these results and suggest avenues for future research.

## 1. Introduction

In 2005, the German Research Foundation (DFG) established the so-called “excellence initiative” (named “initiative” throughout the paper) that ran until 2017. The purpose of the “initiative” was to increase international competitiveness of scientific research at German universities. It involved two funding phases, one in 2006–2011 and one covering 2012–2017. Previous analyses on the “initiative” have mainly examined variables at the university level (“elite universities”), whereas here we home in on the subject level, i.e. academic fields within universities (“university subjects”). In this paper, we describe our investigation of the selection of subject fields within the “excellence initiative,” addressing three interrelated questions. First, which institutional factors mattered most in the selection of subject fields, i.e., why were some fields (in some universities) selected but others (either in the same university or other universities) were not? Second, did the influence of these institutional factors change between the first and second funding phases? Third, to what extent did funding success in the first phase influence funding success in the second phase?

The three research questions have not been empirically examined in the literature on Germany’s “excellence initiative.” Early studies (from the mid-2000s and early 2010s) have shown that funding for the “initiative” was channeled into the largest rather than the most productive universities [1, 2], leading to an increased concentration at larger universities in the system [3, 4]. In addition, some critics argued that universities with high shares of staff in the natural and engineering sciences and with many DFG peer reviewers among their scientific staff received above-average amounts of funding [5–7]. More recent studies (from the mid-2010s and early 2020s) have examined the consequences of the “initiative” for productivity, efficiency, and scientific impact, mostly at the university level [8–17]. Here, we draw on the early studies to formulate hypotheses about selection factors for the “initiative” (Section 2) and discuss empirical results with reference to more recent findings (Section 5).

We argue that examining the *field level* addresses not only an important research gap but also is appropriate for two methodological reasons. First, most of the funding for the “initiative” was channeled into subject fields (3292.2 million Euro, or 71%) via grants for either *Graduate Research Schools* (GS) or *Excellence Clusters* (ECs), and less funding went to *Institutional Strategies* (IS) at the university level (1347.1 million Euro, or 29%) [18]. Thus, the “initiative” primarily funded subject fields rather than entire universities. Second, despite public sector reforms in the 2000s [19–21], German universities have limited strategic planning capabilities compared with higher education institutions in other countries, most notably in the United States [22–26]. Several studies have shown that where planning capabilities have been established at the university level via effective structural reforms, as in the Netherlands in the 1980s and 1990s, universities have achieved consistently high scores in international comparisons of research impact and prestige [27, 28]. In contrast, where such capabilities are less developed, as in German public universities, empirical analyses are better focused on the *field level* [29].

The “initiative” is part of a larger European effort to support global competitiveness of universities via *Centers of Excellence* (CoEs). Among the earliest such initiatives were those in Finland (1995), Norway (2001), and Ireland (2003) [30, 31], and by the mid-2010s, many European countries had established programs, although with considerable variation in goals and funding levels [32, 33]. A review of funding instruments indicates that the size of research grants has increased in most developed countries, and CoE schemes have contributed to this trend [34].

Our analysis yields three empirical insights. *First*, we find that the “initiative” has revealed a stable sorting of public universities into three size classes: small (without “initiative” funding), medium (one or two funding lines), and large (all three funding lines). Our analysis suggests that this sorting already existed in the 1990s and that the “initiative’s” selection procedures have reproduced this size-based order. *Second*, we show that the “initiative’s” selection in the first phase was biased toward large subject fields with considerable grant funding. *Third*, subject fields that received “initiative” support in the first phase were likely to get follow-up support in the second phase, pointing to a high level of path dependence at the field level.

The remainder of this paper is organized as follows: First, we introduce hypotheses based on the available literature (Section 2). We then present the methodology of the paper, including variables and data sources (Section 3). The empirical results follow and include both descriptive statistics and logistic regressions on success in the “initiative’s” two phases (Section 4). Finally, we discuss our findings in light of the recent CoE literature (Section 5).

## 2. Literature review and hypotheses

The “initiative” has attracted considerable attention from academic researchers and political commentators, particularly in the years after it started [35–39], but also more recently [8–17]. Most studies have looked into the “initiative’s” consequences on the public university system, with almost exclusive focus either on the university level (“elite universities”) or the entire higher education system level (“university system”). Some noteworthy results emerged from bibliometric studies; for example, an examination of highly-cited publications (top 10% most cited) found that “the vast majority of universities with Excellence funding held leading or average positions before the funding began. The German university system was already differentiated into stronger and weaker research universities prior to the Excellence Initiative” [12, p. 2234]. In addition, among those universities that published highly-cited research, very few moved upward or downward in the ranking, pointing to a stable sorting within the German university system [11, 13]. Other bibliometric studies reported an increase in productivity but a decrease in scientific impact [17], and an overall loss of efficiency in teaching and research for excellence-funded universities [15, 40]. Even more interesting are results that contradicted earlier warnings about increased concentration of grant funding due to the “initiative”. While both critics and DFG’s leadership envisaged an increased functional differentiation between research universities and teaching colleges [4, 41], such effects have not taken place: Universities with no excellence-funding managed to obtain funding from other sources than the DFG (most notably ministries, and thus other public moneys), thereby both buffering the “initiative’s” effect for individual universities and increasing the level of grant funding in the university system as a whole [8, 16].

While most studies on the “initiative” have examined its consequences on either the organizational or system levels (or both), very few analyses have looked into how its selection procedure functioned at the subject field level. Yet, the early and mostly critical literature on the “initiative” [1–4, 6] is highly informative in that it focused on two arguments, formulated for “elite universities” and the “university system” levels, both of which can be applied to the subject field level as well.

The first argument relates to what Robert K. Merton called the “Matthew effect” [42], a social mechanism that increases inequality (“to those who have, more shall be given”). In the university system, the inequality is between resource-poor and resource-rich public universities, and in brief, the argument is that universities with a large scientific workforce and a high number of research grants are the main beneficiaries of the “initiative” [1, 2, 4, 6, 43]. In contrast, funding is not channeled to the most productive universities, as measured in terms of most publications relative to number of scientific staff [1, 2]. In consequence, absolute size dominates relative performance, contradicting the principle of meritocracy that the “initiative” purports to follow. Based on this argument, which has been discussed almost exclusively at the university level (“elite universities”) and the higher education system level (“university system”), we formulate two hypotheses for the subject/academic field level:

*H1: The probability that a subject field (within a university) will receive excellence funding increases with the absolute size of the field’s scientific staff*.

*H2: The probability that a subject field (within a university) will receive excellence funding increases with the absolute amount of external grant funding to the field*.

A second argument centers on the social construction of scientific (and institutional) prestige. In a more general sense, prestige implies that the production and maintenance of status goods are marked more by notions of quality and refinement than by considerations of utility [44, 45]. In a more specific sense, the early critical commentators argued that rather than simply identifying and awarding support for truly excellent research, the “initiative” was involved in an “act of consecration”, a procedure that bestows scientific prestige on universities despite apparent inefficiencies and low relative performance. This “consecration” was typically performed by professors from the resource-rich universities (partly from abroad, but also from Germany) and follows the logic of supporting those who have already accumulated considerable resources, thus deepening the consequences of the Matthew effect [1, 7].

Based on this claim, we probe whether selection decisions in the first funding phase consecrated those chosen to receive the funding, and in this way helped them obtain funding in the second phase, as well. So far, the scientific prestige argument has been predicated on findings at the university level, but it can be extended to the level of academic fields [46, 47]. Hence, we formulate the following hypothesis:

*H3: The probability that a subject field (within a university) will receive excellence funding in the second phase increases when it has been designated as excellent in the first phase*.

## 3. Methodology

### 3.1 Variables

Our dependent variable (DV) is binary and measures whether or not a *university subject* received “initiative” funding (via either the GS or the EC funding line or both) between 2006 and 2011 (first phase) and/or between 2012 and 2017 (second phase).^1^ Our DV differs from the early literature [1, 2, 6] and some of the later efficiency-oriented literature [15, 17, 40] that centered around the amount of grant funding, highly-cited publications and citations, each per professor or scientific staff (a proxy for research efficiency). We use a binary DV for two reasons. First, the DFG does not provide the amounts of funding per funding line and subject field in a given university. Therefore, a cardinal ranking using “initiative” funding at the field level within universities is simply not available. Second, we are interested in estimating the chances of subject fields within universities (“university subjects”) to receive “initiative” funding based on explanatory variables (see below). Therefore, information about whether or not (1/0) a subject field in a given university has received “initiative” funding provides sufficient information for carrying out this estimation procedure (non-linear probabilistic regression, here: logit regression).

Our independent variables (IVs) are measured at the level of university subjects *(field level)* and include number of professors (IV1), amount of all grant funding in million Euros (IV2), and a binary variable for the second phase that measures whether a university subject was funded in the first phase (IV3). Another IV is measured at the *university level*: amount of DFG grant funding in million Euros (IV4). It should be pointed out that the “initiative’s” focus is on research, and not teaching. Therefore, all IVs – as explanatory variables – are meant to capture various aspects of the research dimension. Professors (IV1) are the most senior level of scientists recruited for doing research (and teaching), and grant funding (IV2, IV4) measures the amount of externally funded research activities both guided (and carried out) by professors. In addition, IV3 records the continuity of “initiative” funding in both the first and second phase.

In addition, we included one control variable (CV), the number of students (CV1). CV1 was entered to control for the size of a subject field’s student population. While the IVs are theoretically anchored (Matthew effect, prestige consecration), the CV is introduced to reduce errors in measurement. Initially, the number of non-professorial scientific staff, amount of basic funding in million Euros, number of WoS publications, and number of WoS citations were included in our list of CVs. However, because these four variables were highly correlated with IV1 and with each other, we excluded them from further analyses to avoid multicollinearity (Appendix 7). For robustness checks, we also conducted a regression analysis using the number of citations as a control variable (to account for subject field differences in scientific visibility) instead of DFG grant funding. We made this change because both variables are highly correlated. However, this change did not significantly alter the main results (see Appendix 11).

**Table 1:** List of dependent, independent and control variables.

| **Dependent variable** |  |
| --- | --- |
| DV: Excellence initiative funding | Binary (1/0) |
| **Independent variables** |  |
| IV1: Number of professors (field level) | Total number |
| IV2: Total grant funding (field level) | Million Euros |
| IV3: Funded in first phase (field level) | Binary (1/0) |
| IV4: DFG grant funding (university level) | Million Euros |
| **Control variable** |  |
| CV1: Number of students (field level) | Total number |

Regarding H1, IV1 was used to operationalize the absolute size of (senior) scientific staff. With respect to H2, IV2 and IV4 were used to capture the absolute amounts of grant funding, both at the subject field level (IV2) and the university level (IV4). With regard to H3, we used IV3 to examine the effects of prior funding success on the subsequent probability of receiving “initiative” funding. All IVs and CV were measured in the five years prior to the first phase (beginning in 2006) and the second phase (beginning in 2012). Therefore, we computed mean values for IV1–4 and CV1 for the years 2001–2005 and then for the years 2007–2011. As a robustness check, we also computed mean values for the three years preceding the first and second phases (i.e., 2003–2005 and 2009–2011). We found no major differences between the different time windows, so our interpretation builds on the five-year time windows.

Due to the nature of our DV, we calculated logistic regression models to measure the effects of the IVs on the odds of receiving “initiative” funding [48, 49]. The logistic regression is the preferred method due to the binary measurement of our DV. To ensure that our models fit appropriately, we calculated goodness of fit tests (Appendix 10) developed by Hosmer, Lemeshow (50).

In addition to the results presented below (Section 4), we calculated two-level regression models with subject fields as the first-level unit and universities as the second-level unit. As mentioned above (Section 1), most empirical studies have focused on variables at the university level (“elite universities”), so we added two time-invariant binary variables to the existing IVs and CV at the university level: one that measures whether universities were founded after 1945 (“young” universities) and one that captures the geographical location (“West Germany” versus “East Germany”). With these additional models, we controlled for variance at the university level [51] (Appendix 9).

### 3.2 Units of Analysis

This paper focuses on public universities in Germany. According to the Federal Statistical Office (StBA) in 2015, of 102 universities with permission to award doctoral degrees, 82 are state-run and 20 are privately run. Private universities play only a minor role due to their low share of all enrolled students, the very limited range of disciplinary fields offered, and the low level of research activities [19]. They are therefore not considered further here. In addition, we excluded several specialized public higher education institutions that are not suitable for comparison with universities that offer a broad range of fields; and some universities could not be analyzed due to considerable gaps in the data, especially staffing and funding data (for more details, see [26]). Of the remaining universities, 17 were classified as “technical” (TUs, Appendix 1), either because they are members of the TU9 association of the nine leading technical universities in Germany or because they bear “Technische Universität” or “Technische Hochschule” in their name; the remaining 51 universities were classified as “non-technical” (NTUs, Appendix 2).

Data on staff, finances, and student numbers for these 68 public universities (1995–2018) were obtained directly from the StBA (Appendix 3) and correspond with published data from the reports in Serial 11: Education and Culture, Sections 4.1 (Students at Universities), 4.4 (Personnel at Universities), and 4.5 (Finances at Universities). For students, the StBA records the target degrees grouped into accumulated higher-level categories, and of these, here we use university degrees, bachelor’s and master’s degrees, and teaching examinations (Appendix 4). Medicine was excluded because separating hospital units from their affiliated university departments was not possible for all years.

Publication and citation data for the 68 universities were obtained from the Competence Centre for Bibliometrics, using the WoS. From this source, cleaned address data are available at the university level but not at a more detailed level of aggregation, which means publications cannot be directly allocated to institutes, academic departments, or faculties. Instead, we used a field classification to break down universities into their subject fields. Following [26], we use the Archambault classification [52], which assigns each journal to one subject category and provides a good match with the StBA field classification (Appendix 5). In this way, we connected WoS publication and citation data with the staff, finance, and student data provided by the StBA.

For our purposes, each WoS publication was counted once, even if several co-authors from a single university were listed. The procedure was different for publications with co-authors from different institutions; in this case, the WoS publication was counted for each institution (whole count). We considered all publications in journals covered by the WoS with the participation of at least one German university that belonged to the document type “Article,” “Review,” or “Letter.” The same types of documents were considered for measuring citations. We counted all citations received by a university subject per year, regardless of the publication years of the cited publications. With this counting method, the change in the citations obtained for a subject field in the respective year could easily be observed.

The validity of bibliometric analyses depends on the rate of coverage for the respective subject field. To estimate the coverage for German publications by subject fields, we analyzed the proportion of their cited references that were in turn included in the WoS. This method is called “internal” WoS field coverage [53] and provides a proxy for how well the WoS reflects scholarly activity of an academic field. Our analysis documented that the internal coverage in many fields is insufficient for bibliometric analyses. Following a common standard [53], we applied a cut-off value of 50% for cited references. This cut-off yielded the following subject fields: Biology, Chemistry, Physics and Astronomy, Psychology, Agricultural Sciences, Food and Beverages Technology, Mechanical Engineering/Process Engineering, Geosciences [excluding Geography], Electrical Engineering, Forestry/Timber Management, Economics, and Mathematics (Appendix 5, see also Fig. 1 in [26]). For these 12 subject fields, we obtained and analyzed publication and citation data.

Furthermore, we retrieved data from reports on the distribution of “initiative” funding [18, 54, 55]. We began with detailed descriptions of GS and ECs of the second funding phase. In these cases, the official spokesperson for each project is named, allowing for a clear assignment to a particular subject field. In addition, the descriptions often contain the affiliation of the project to one or several subject fields. Based on the official spokesperson and the GS or EC descriptions, we were able to assign them to subject fields using the StBA classification. Every GS or EC was assigned to at least one – and up to three – such subject fields.

The assignment of funded projects from the first phase required more work. Of 83 funded GS or ECs, 70 were continued in the second phase, so that only 12 were not yet assigned to a subject field. The DFG assigns each GS or EC to one disciplinary domain: humanities and social sciences, life sciences, natural sciences, or engineering [54]. We differentiated these into subject fields, either directly through the name (e.g., “Bonn Graduate School of Economics” was assigned to economics) or if an assignment was not possible, by conducting searches of other (internet) sources for official spokespersons, publications, or public events that allowed an assignment to up to three disciplines. This methodology produced Appendix 6, which lists all three lines of “initiative” funding at our 68 universities. We dispensed with a verbatim catalogue, which would have been untenably extensive.

All data were processed at the level of universities and their respective subject fields. Here, we use the term “university subject” to denote *subject fields within universities*. With 68 universities and 56 subject fields, we arrived at 68 * 56 = 3,808 possible observations with respect to staff, funding, and student data, of which, for example, 2,829 were realized for grant funding. For our analyses, we reduced the sample size to only valid observations in all of our variables of interest for the relevant years. Regarding the bibliometric analyses, the number of fields is 12, so that we arrive at 68 * 12 = 816 possible observations, of which, for example, 624 were realized for publications.

## 4. Analysis and Discussion of Results

### 4.1 Descriptive Results

A first set of descriptive results provides contextual information at the university level for H1 and H2. Starting with an overview of changes across *all universities* from 1995 to 2018, there were considerable differences with regard to average growth between the IVs and CVs during the observation period (Table 2, see “mean values”). The average number of professors (IV1) grew by 8% (from 254.7 to 276.0), the amount of DFG grant funding at the university level (IV4) by 215% (from 10.4 million Euros to 32.9 million Euros), and total grant funding at the subject field level (IV2) by 150% (from 23.8 million Euros to 59.5 million Euros); the number of WoS citations skyrocketed by 886% (from 4,812 to 47,440) and the number of students (CV1) by 26% (from 15,902 to 19,997). These results suggest that German professors not only taught more students in 2018 (compared with 1995) but also that their research activities, as exemplified by external grant moneys, increased over time.

**Table 2:** Variables by “excellence initiative” funding status.

|  | Mean values |  |  |  |  |  | Factor values |  |  |  |  |  |
| --- | --- | --- | --- | --- | --- | --- | --- | --- | --- | --- | --- | --- |
|  | 1995 | 2000 | 2005 | 2010 | 2015 | 2018 | 1995 | 2000 | 2005 | 2010 | 2015 | 2018 |
| **Number of professors** |  |  |  |  |  |  |  |  |  |  |  |  |
| All universities | 254.7 | 247.7 | 240.0 | 256.3 | 274.1 | 276.0 | 1.00 | 1.00 | 1.00 | 1.00 | 1.00 | 1.00 |
| Non-funded universities | 174.3 | 183.8 | 177.0 | 178.5 | 189.6 | 187.2 | *0.68* | *0.74* | *0.74* | *0.70* | *0.69* | *0.68* |
| Universities with at least one funding line | 304.5 | 287.2 | 279.0 | 304.4 | 326.4 | 331.0 | *1.20* | *1.16* | *1.16* | *1.19* | *1.19* | *1.20* |
| Universities with all three funding lines | 361.1 | 336.7 | 326.6 | 357.6 | 389.9 | 400.6 | *1.42* | *1.36* | *1.36* | *1.40* | *1.42* | *1.45* |
| **Total grant funding** |  |  |  |  |  |  |  |  |  |  |  |  |
| All universities | 23.8 | 28.9 | 28.8 | 48.8 | 54.8 | 59.5 | 1.00 | 1.00 | 1.00 | 1.00 | 1.00 | 1.00 |
| Non-funded universities | 12.9 | 14.6 | 14.3 | 24.6 | 28.0 | 28.5 | *0.54* | *0.51* | *0.50* | *0.50* | *0.51* | *0.48* |
| Universities with at least one funding line | 30.6 | 37.7 | 37.9 | 63.7 | 71.3 | 78.7 | *1.28* | *1.31* | *1.31* | *1.31* | *1.30* | *1.32* |
| Universities with all three funding lines | 43.5 | 55.8 | 57.3 | 91.0 | 100.2 | 115.0 | *1.83* | *1.93* | *1.99* | *1.86* | *1.83* | *1.93* |
| **DFG grant funding** |  |  |  |  |  |  |  |  |  |  |  |  |
| All universities | 10.4 | 13.5 | 13.6 | 23.7 | 27.1 | 32.9 | 1.00 | 1.00 | 1.00 | 1.00 | 1.00 | 1.00 |
| Non-funded universities | 4.4 | 5.8 | 5.5 | 9.1 | 9.0 | 11.0 | *0.42* | *0.43* | *0.41* | *0.38* | *0.33* | *0.34* |
| Universities with at least one funding line | 14.0 | 18.2 | 17.8 | 32.8 | 38.3 | 46.4 | *1.34* | *1.35* | *1.31* | *1.38* | *1.41* | *1.41* |
| Universities with all three funding lines | 20.7 | 24.2 | 22.9 | 47.4 | 56.4 | 68.3 | *1.98* | *1.80* | *1.69* | *2.00* | *2.08* | *2.08* |
| **WoS citations** |  |  |  |  |  |  |  |  |  |  |  |  |
| All universities | 4.8 | 8.3 | 12.9 | 21.1 | 34.3 | 47.4 | 1.00 | 1.00 | 1.00 | 1.00 | 1.00 | 1.00 |
| Non-funded universities | 1.5 | 2.8 | 4.6 | 7.8 | 12.9 | 18.4 | *0.32* | *0.34* | *0.36* | *0.37* | *0.38* | *0.39* |
| Universities with at least one funding line | 6.8 | 11.6 | 18.0 | 29.3 | 47.6 | 65.4 | *1.42* | *1.41* | *1.40* | *1.39* | *1.39* | *1.38* |
| Universities with all three funding lines | 9.6 | 16.1 | 24.5 | 40.6 | 67.5 | 94.3 | *2.00* | *1.95* | *1.90* | *1.93* | *1.97* | *1.99* |
| **Number of students** |  |  |  |  |  |  |  |  |  |  |  |  |
| All universities | 15.9 | 15.0 | 15.9 | 16.2 | 19.6 | 20.0 | 1.00 | 1.00 | 1.00 | 1.00 | 1.00 | 1.00 |
| Non-funded universities | 9.2 | 9.4 | 10.6 | 11.5 | 13.5 | 13.9 | *0.58* | *0.63* | *0.66* | *0.71* | *0.69* | *0.69* |
| Universities with at least one funding line | 20.1 | 18.4 | 19.2 | 19.1 | 23.4 | 23.8 | *1.26* | *1.23* | *1.21* | *1.18* | *1.19* | *1.19* |
| Universities with all three funding lines | 24.0 | 21.6 | 22.6 | 21.7 | 26.5 | 26.7 | *1.51* | *1.44* | *1.42* | *1.34* | *1.35* | *1.33* |

Notes: funding numbers are in million Euros (inflation adjusted, base: 2005), student numbers and WoS citations in thousands; grand funding represents income (not spending). Factor values = mean values, normalized with all universities. Variables in this table are measured on the university level. “DFG grant funding” variable is identical to IV4 (university level).

We probed whether universities without any “initiative” funding differed in their growth pattern compared with universities with some or full “initiative” funding. For this purpose, we mapped all IVs and CVs onto three university categories, beginning in 1995 and with the university system as reference category (Table 2). We note that the three university categories were devised using information on both the first and second phases of the “initiative”: (1) “non-funded universities” were those with no funding in both the first and second phases, (2) “universities with at least one funding line” included all universities that received either GS or EC in the first and/or second phase(s), and (3) “universities with all three funding lines” included universities that received all three lines of funding either in the first and/or second phase(s).

Based on this tabulation, we observed a very stable social order within the German university system that appeared to be based on organizational size: <u>small</u>, <u>medium</u>, and <u>large universities</u> (Table 2). This means that the number of funding lines received by a university increases with the size of the university, which is reflected in the number of scientific staff. It is important to emphasize that this size-based order already existed in the 1990s: There was almost no change between 1995, 2000, and 2005 – the years before the “initiative” started to distribute funding; in addition, there was very little change between 2010, 2015, and 2018.

The size-based social order of public universities in Germany means that universities with no “initiative” funding represent, on average, the smallest entities, followed by medium-sized institutions that received support from at least one line of funding (GS or EC) and the largest ones that received support from all three lines of funding (GS, EC, IS). To illustrate the changes relative to the mean values in each year, we calculated factor scores to show whether universities with different funding lines were below or above average in terms of staff, grants, citations, and students (Table 2, see “factor values”). Regarding professorial staff, non-funded universities were about 70% of the average size in 1995; regarding total grant funding, they reached about 50% of the mean; and regarding citations, they had about a third of the mean number of citations. Universities with at least one funding line were bigger and had greater scientific visibility: Their professoriate was 20% above the average, their total grant funding was 30% above average, and their citations were 40% above the means. The “excellence” universities were the largest institutions: Their professoriate was 40% and their total grant funding 80% above the mean, and they had about twice as many citations as the average university. The magnitude of these ratios has remained robust over time.

The descriptive results are interesting with regard to H1 and H2 in that large universities with above-average amounts of external grant funding in the years prior to the “initiative” are those that received most “excellence” funding. As shown below, these findings are supported by logistic regressions at the subject field level as well. In addition, the results are consistent with regard to scientific visibility. Universities with above-average numbers of citations in the years preceding the “initiative” were those that received the most “initiative” funding.

Regarding student numbers, “excellence” universities had, on average, about twice as many students as non-funded institutions in 2018, yet they were larger in relative terms 23 years earlier when they had about three times as many students (compared with non-funded universities). More specifically, “excellence” universities had about 24,005 students in 1995, compared with 26,690 in 2018 (plus 11%), whereas non-funded universities had about 9,174 students in 1995 and 13,854 in 2018 (a 51% increase). In other words, the overall growth of students between 1995 and 2015 was absorbed (both in absolute and relative terms) to a greater extent by universities with no “initiative” funding compared with those that received full funding.

A second set of descriptive results provides contextual information for H3 at both the subject field level and the university level between 2006 and 2017 (Table 3a). A cross-tabulation between subject fields that received “initiative” funding in either one (GS or EC) or two lines (GS and EC) in the two phases revealed a staggering level of path dependency. We observe 2,248 subject fields in the first phase that received no funding, of which 2,215 (99%) received no funding in the second phase either. Among the 102 subject fields with one or two lines of “initiative” funding in the first phase, 91 (89%) received funding in the second phase as well. The overall “reproduction rate” at the subject field level, i.e., the diagonal of Table 3a, divided by all observations, amounted to 98% (2,300/2,350). Based on a chi-square test, independence between the two funding phases was rejected at a high level of statistical significance.^2^

**Table.3a:** Number of subject fields by funding phase.

|  | 2012–2017 |  |  |  |
| --- | --- | --- | --- | --- |
| 2006–2011 | No funding | One line | Two lines | Total |
| No funding | 2,215 | 33 | 0 | 2,248 |
| One line | 11 | 77 | 4 | 92 |
| Two lines | 0 | 2 | 8 | 10 |
| Total | 2,226 | 112 | 12 | 2,350 |

Chi<sup>2</sup> = 2600 // $p = 0.000$
Notes: one line = GS or EC; two lines = GS and EC.

While Table 3a provides information at the subject level, Table 3b offers funding information at the university level. We observed 35 universities with no “initiative” funding in the first phase, meaning that none of their subject fields received any “initiative” funding. Of these, 27 (77%) also received no follow-up funding in the second phase (Table 3b). Among the 33 universities with at least one line of “initiative” funding in the first phase, 32 (97%) also received such funding in the second phase. The “reproduction rate,” i.e., the diagonal in Table 3b, divided by all observations, amounts to 72% (49/68). Again, based on a chi-square test, independence was rejected at a high level of statistical significance.

**Table 3b:** Number of universities by funding phase.

|  | 2012–2017 |  |  |  |  |
| --- | --- | --- | --- | --- | --- |
| 2006–2011 | No funding | One line | Two lines | Three lines | Total |
| No funding | 27 | 8 | 0 | 0 | 35 |
| One line | 1 | 10 | 1 | 1 | 13 |
| Two lines | 0 | 1 | 7 | 4 | 12 |
| Three lines | 0 | 1 | 2 | 5 | 8 |
| Total | 28 | 20 | 10 | 10 | 68 |

$\chi^2 = 78.3793 // p = 0.000$
Notes: one line = GS or EC; two lines = GS and EC; three lines = GS and EC and IS.

Taken together, these results and in particular those from Table 3a provide support for H3: Subject fields (and universities) with funding in the “initiative’s” first phase were extremely likely also to receive support in the second phase, a clear indication of path dependency. As shown below, logistic regressions provide additional support for this descriptive finding.

### 4.2 Results from logistic regressions

We conducted separate logistic regression analyses for each funding phase, as outlined above. Our results, using observation of all universities and thus our largest sample, provided considerable support for H1–H3. There were some differences and modifications with regard to NTUs and TUs on the one hand and subject fields with good bibliometric coverage on the other hand (see Appendix tables). In addition, we follow Breen et al.’s recommendation [49] that the absolute magnitude of IVs’ coefficients should not be compared across models, but rather their direction (either positive or negative) and statistical significance (p-value).

*First*, and in support of H1, our analysis with all universities (Table 4a, b) revealed that in both funding phases, the number of professors significantly increased the chances for a subject field to be selected by the “excellence initiative” (all models). This result confirms similar findings at the university level [8]. *Second*, and in support of H2, the amount of external grant funding had explanatory power as well. In the first phase, both total grant funding at the subject field level (IV2) and DFG funding at the university level (IV4) significantly reduced the share of unexplained variance in the dataset (Table 4a), and in the second phase, these two variables were significant (model 4; Table 4b) only before IV3 was introduced (model 5; Table 4b). *Third*, and in strong support of H3, success in the second phase of the “initiative” largely depended on success in the first phase.

**Table 4a:** Logistic regression, first “initiative” phase (2006–2011), all universities.

|  | DV = Excellence Initiative funded |  |  |  |
| --- | --- | --- | --- | --- |
|  | Model 1 | Model 2 | Model 3 | Model 4 |
| Professors | 0.105099*** | 0.079204*** | 0.075300*** | 0.087418*** |
| Total grant funding |  | 0.130803*** | 0.097092** | 0.089602** |
| DFG grant funding |  |  | 0.057499*** | 0.058350*** |
| Students |  |  |  | -0.196914 |
| Intercept | -4.281834*** | -4.223346*** | -5.131329*** | -5.138308*** |
| Observations | 2,388 | 2,388 | 2,388 | 2,388 |
| Pseudo-R2 | 0.152671 | 0.173839 | 0.201265 | 0.203428 |

\* $p < 0.05$ , \*\* $p < 0.01$ , \*\*\* $p < 0.001$ ; ExIn = “excellence initiative”

**Table 4b:**
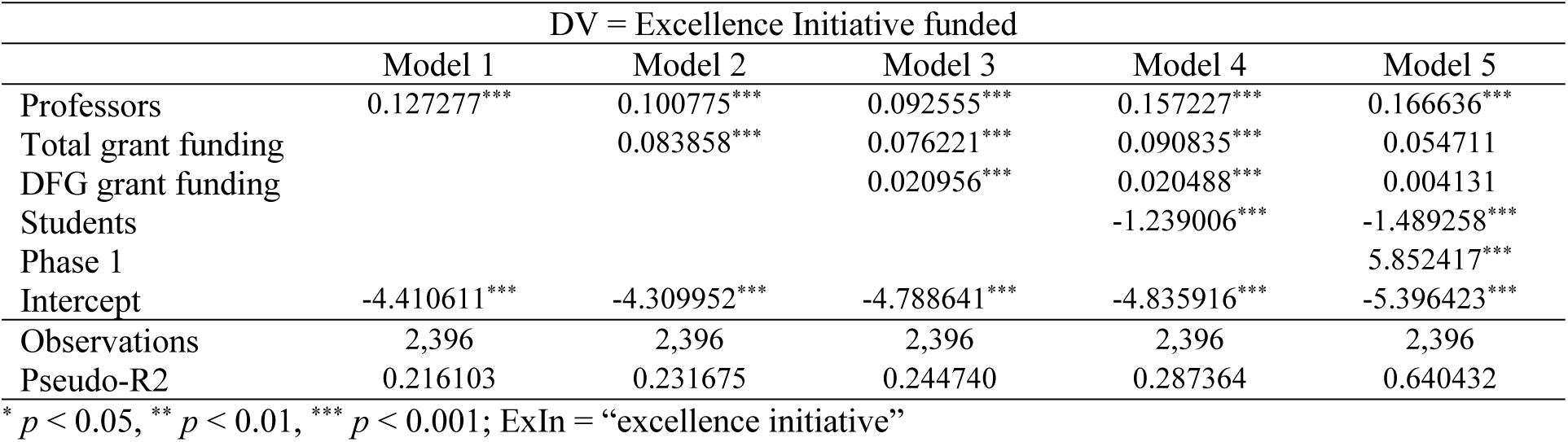
Logistic regression, second “initiative” phase (2012–2017), all universities.

There were some noteworthy differences regarding university groups. For NTUs, grant funding was highly significant in the second phase as well, even when the consecration variable (IV3) was introduced, suggesting that grant funding was a more decisive factor in selecting subject fields at NTUs compared with TUs (Appendix Table 8a, 8b). In contrast, for TUs (Appendix Table 8b) and similar to the main results across all universities (Table 4b), grant funding is not a significant predictor of excellence funding in the second phase if they received excellence funding already in the first phase (IV3).

Broadly defined academic domains also differed. Although results for the natural sciences (Appendix Table 8e-f) were quite similar to the analyses with all universities presented above, the humanities and social sciences were less complex. For the latter, grant funding (either IV2 or IV4) in the first “initiative” phase and prestige (IV3) in the second “initiative” phase were the only significant factors (Appendix Table 8g-j). Furthermore, when we examined subject fields with good WoS coverage, scientific visibility emerged as a relevant factor, in addition to grant funding (first phase), professors (second phase), and prestige (both phases) (Appendix Table 8k-l).

We also note that adding the university level to the logistic regression analyses did not add explanatory power to our models. Neither foundation year nor geographical location had significant effects in our two-level analyses (Appendix 9).

Taken together, these findings are largely consistent with the descriptive results: The more professorial staff (IV1) and the more external grant funding (IV2) associated with a subject field, the more likely it was to obtain “initiative” funding. Also, the second phase was dominated by the first phase (IV3): Subject fields that received support in the first phase were extremely likely also to do so in the second phase.

Our results provide strong support for the early and mostly critical literature in that absolute size and socially constructed scientific prestige emerged as important institutional factors predicting success in the “initiative”. Although early analyses focused mainly on the university level (“elite universities”), we show that size and prestige also can be found at the more disaggregated level of subject fields.

## 5. Conclusion

The fact that subject fields flush with research cash had better chances of securing “initiative” support points to *selection procedures that were biased towards grants*. Gerhards [43] argues that grant funding seems to be a “fetish” in the German higher education landscape, particularly with regard to the “initiative”: Although it should be regarded simply as an input for conducting research, organizational actors in German science policy, most notably the DFG and the German Science and Humanities Council (Wissenschaftsrat) [54, 56], have cultivated the notion that grant funding is equivalent to research success. Gerhards writes (our translation): “It produces a biased picture when the DFG publishes mostly absolute measures which are not weighted by staff. Yet, this biased picture has been taken up gratefully by large institutions which use it for self-staging and self-advertising” [43, pp. 44-46]. Gerhards goes on to say that using grant funding as a proxy for research success would be legitimate only if grants and publications were highly correlated. However, empirical evidence suggests otherwise, and he recommends that “a better institutionalization of bibliometric methods in Germany would align incentives with international standards and, in the long run, could improve Germany’s research performance in international comparisons” [43, p. 50]. We will return to this latter point below.

We found that subject fields with many professors and thus *large disciplinary entities were more successful in getting “initiative” support than smaller ones*. This result contrasts to a growing body of literature showing that breakthrough science is associated with small research entities, whereas large teams typically develop and exploit existing scientific programs [57–59]. Scientific excellence seems to be not so much a feature of “big science” but rather typical of “little science,” to borrow a book title from de Solla Price [60]. In addition, evidence from the Nordic countries suggests that large grants given to Centers of Excellence (CoE) did not have the highest impact when they were awarded to already highly ranked research groups but rather when they were “awarded to groups not yet performing at the highest level” [61]. Similarly, a large-scale bibliometric study from Canada shows “that in terms of both the quantity of papers produced and their scientific impact, the concentration of research funding in the hands of the so-called ‘elite’ of researchers generally produces diminishing marginal returns” [62]. These findings are corroborated by a recent review that finds, as a general result, “diminishing returns to grant size, measured for example in terms of number of publications, citation impact and number of highly cited papers” [34].

Furthermore, only between 102 (first phase) and 124 (second phase) subject fields among 2,350 examined (4.3% and 5.3%, respectively) were funded under “initiative’s” umbrella. This suggests that the *“initiative’s” potential impact on improving research capacities must have been very limited*. Bonaccorsi et al. [28] showed that 16 German universities with 34 subject fields produce research that scores among the 10% most cited publications worldwide, a rather small number of entities compared with the Netherlands, which scores with 12 universities and 37 subject fields. It seems plausible to assume that the “initiative’s” impact would have been greater had it provided funding for a substantially larger number of subject fields. Of course, such an impact would have been possible only with a much larger investment in universities’ infrastructures [63, 64].

Finally, there is the question of whether the “initiative” had any impacts so far, in particular with regard to its institutional mission of supporting “research excellence.” In our view, although the Matthew effect in science has gained renewed attention in recent years, with several studies providing evidence for its continued relevance [65–68], available empirical studies suggest that this social mechanism was less important for the “initiative” than some had initially thought. *In financial terms, the “initiative” has not increased inequality between funded and non-funded universities* so far. There has been an overall upward shift to a higher funding level for all universities [16] because those without “initiative” funding managed to find other (mostly public) sponsors [8]. These results are in line with our descriptive finding that Germany’s public university system was stable between 1995 and 2018 (Section 4.1).

*In terms of research impact, current evidence suggests no major changes resulting from the “initiative”.* Based on bibliometric findings [12, 13], the initiative’s international evaluation commission concluded that although “bibliometric investigations show an impressive qualitative performance regarding publications stemming from Excellence Clusters,” it remains “unclear to what extent new research priority areas emerged due to the support from the Excellence Initiative or whether the Excellence Initiative has instead led to a bundling of existing research capacities and hence increased visibility” [69, p. 5]. Recent bibliometric evidence suggests that universities funded through the “initiative” have shown a decreasing citation impact, whereas universities without such support have increased their citation rates [10, 17].

Perhaps most importantly, early commentators believed that there would be an increasing functional differentiation between teaching and research universities. For example, Hartmann argued (our translation): “The German university system faces a permanent split between two types of universities: research universities and vocational universities. Research will be concentrated at the former, while the latter will conduct almost no research (like today the universities of applied sciences) but quickly prepare students for their job” [4]. Similarly, Winnacker, president of the DFG during implementation of the “initiative’s” first funding phase, emphasized the increasing functional differentiation (our translation): “The differences in quality between the universities are already considerable, they will grow further through the Excellence Initiative. (…) The [university] system will differentiate further. In addition to pure research universities that will follow standards of modern scientific research in their education, there will be universities that will attempt such standards in a few subject fields only, universities that will not even strive for such standards, and universities that will develop their strengths in practical orientation” [41].

However, *no empirical evidence so far supports the far-reaching claim of an increased functional differentiation between teaching and research universities due to the “initiative”*. Even the international evaluation commission concluded that “it is not possible to demonstrate an increased differentiation of the German university system as a whole as a consequence of the Excellence Initiative” [69, p. 5].

In summary, when we consider that the “initiative’s” selection procedures were biased towards large disciplinary entities with considerable grant money, leading to the continued support of very few subject fields, two tentative policy recommendations seem justified.

*First*, we reiterate Gerhards’ point that the selection of fields should be based on research performance, using bibliometric methods that follow international standards [70]. The current practice of equating grant funding with research quality has clearly not improved the research performance of German universities. Rather, we have reason to believe that the exclusive focus on grant funding has set incentives for research groups to grow, irrespective of their marginal productivity and their capabilities to conduct breakthrough research.

*Second*, the limited reach of support for subject fields (4% of all possible fields) raises the question of the “initiative’s” effectiveness in upscaling internationally competitive research capacities. The current practice of selecting very few subject fields in very few universities has not improved such capacities on a measurable level. To be globally competitive, German universities need much larger long-term financial support; otherwise, they will continue to trail North American and increasingly also Asian universities [28, 64, 71].

Our study has a potential weakness: since we do not have access to the exact amounts of the *“initiative’s”* funding for subject fields in universities (“university subjects”), we were not able to calculate (OLS) regression analyses using a cardinal dependent variable. Therefore, we welcome future studies with more disaggregated funding data, preferably retrieved with additional support from the DFG.

Finally, our analysis covers the first and second phase of the *“initiative”,* yet it is quite plausible that its successor, the “excellence strategy” continues the existing structural pattern. Therefore, we encourage future studies to look into continuities between the “excellence initiative” and the “excellence strategy,” with special focus on whether there has been upward or downward mobility of university subjects (and entire universities).

## Appendices

## Appendix 1: List of technical universities (alphabetical)

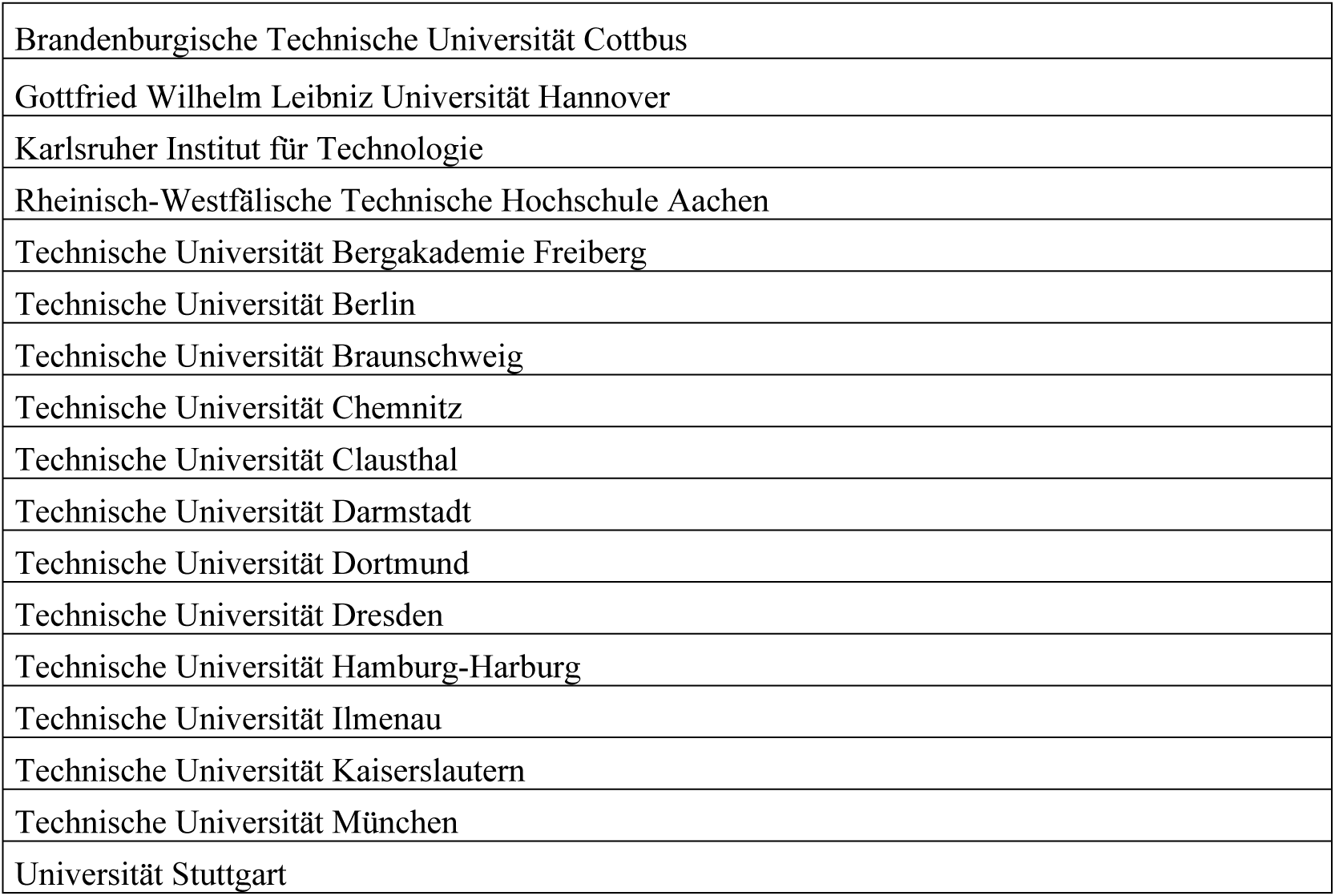

## Appendix 2: List of non-technical universities (alphabetical)

|  |
| --- |
| Albert-Ludwigs-Universität Freiburg |
| Bergische Universität Wuppertal |
| Carl von Ossietzky Universität Oldenburg |
| Christian-Albrechts-Universität zu Kiel |
| Eberhard Karls Universität Tübingen |
| Ernst-Moritz-Arndt-Universität Greifswald |
| Europa-Universität Viadrina Frankfurt (Oder) |
| Freie Universität Berlin |
| Friedrich-Alexander-Universität Erlangen-Nürnberg |
| Friedrich-Schiller-Universität Jena |
| Georg-August-Universität Göttingen |
| Goethe-Universität Frankfurt am Main |
| Heinrich-Heine-Universität Düsseldorf |
| Humboldt-Universität zu Berlin |
| Johannes Gutenberg-Universität Mainz |
| Julius-Maximilians-Universität Würzburg |
| Justus-Liebig-Universität Gießen |
| Ludwig-Maximilians-Universität München |
| Otto-Friedrich-Universität Bamberg |
| Otto-von-Guericke-Universität Magdeburg |
| Philipps-Universität Marburg |
| Rheinische Friedrich-Wilhelms-Universität Bonn |
| Ruprecht-Karls-Universität Heidelberg |
| Universität Augsburg |
| Universität Bayreuth |
| Universität Bielefeld |
| Universität Bochum |
| Universität Bremen |
| Universität des Saarlandes |
| Universität Duisburg-Essen |
| Universität Erfurt |
| Universität Halle |
| Universität Hamburg |
| Universität Hohenheim |
| Universität Kassel |
| Universität Koblenz-Landau |
| Universität Konstanz |
| Universität Leipzig |
| Universität Lüneburg |
| Universität Mannheim |
| Universität Osnabrück |
| Universität Paderborn |
| Universität Passau |
| Universität Potsdam |
| Universität Regensburg |
| Universität Rostock |
| Universität Siegen |
| Universität Trier |
| Universität Ulm |
| Universität zu Köln |
| Westfälische Wilhelms-Universität Münster |

## Appendix 3: Data from the Federal Statistical Office (StBA)

Outliers and missing data in the personnel, financial and student variables were identified and missing values were imputed wherever possible in an otherwise steady trend. Outliers that differed by several orders of magnitude from the preceding and following years were recoded as missing values.

The staff statistics from the Federal Statistical Office identify five non-overlapping categories for professorial staff: “temporary professors,” “C4/W3 professors,” “C3/W2 professors,” “C2 professors,” and “junior professors.” These categories constitute the variable “professors” in our data set.

The financial statistics from the Federal Statistical Office distinguish between two categories of external funding: “public grants” and “grants from other areas,” which together constitute the variable “revenue from grants.” They have been adjusted for inflation (base year: 2010).

The student statistics from the Federal Statistical Office differentiate among registered students according to their study objective (target degree). We have used the available categories to produce the variable “total students,” which comprises students working toward Bachelor’s and Master’s degrees as well as students aiming for a teaching degree.

## Appendix 4: Aggregation scheme of StBA examination groups

|  | **Aggregated group** | **Examination groups (standard font), final examinations (*italic*)** |
| --- | --- | --- |
| X | University degree (without teacher training examinations) | *Master degree; licentiate; ecclesiastical examination; state examination/1<sup>st</sup> state examination; state examination (single-phase education); faculty examination; diploma (U); diploma (U-GH); diploma (U) - interpreter; diploma (U) - translator; certified translator; diploma (U) - teacher; final examination without academic degree* |
| X | Bachelor degree (without teacher training examinations) | *Multi-field bachelor without teacher training option; multi-field bachelor with teacher training option; bachelor (U) - teacher; bachelor at universities; bachelor at art colleges; bachelor at universities of applied sciences* |
| X | Master degree (without teacher training examinations) | *Multi-field master; master (U) – teacher; master at universities (final examination required); master at art colleges (final examination required); master at universities of applied sciences (final examination required)* |
| X | Teacher training examinations | Teacher training, bachelor and master examinations (TT/BA/MA) at primary and lower secondary schools/primary level; teacher training, bachelor and master examinations (TT/BA/MA) at intermediate level (secondary level I/primary schools/primary level); teacher training, bachelor and master examinations (TT/BA/MA) at secondary schools/technical secondary level I; teacher training, bachelor and master examinations (TT/BA/MA) comprehensive examination for secondary level II/secondary level I; teacher training, bachelor and master examinations (TT/BA/MA) at grammar school/secondary level II, general schools; teacher training, bachelor and master examinations (TT/BA/MA) at special schools; teacher training, bachelor and master examinations (TT/BA/MA) at vocational schools/secondary level II, vocational schools; TT bachelor; TT master; other TT according to school types/school levels |
|  | Doctorates | Doctorates |
|  | Artistic degree | Artistic degree (not including *bachelor degree at art colleges* or *master degree at art colleges (final examination required)*) |
|  | University of Applied Sciences degree | University of Applied Sciences degree (not including *bachelor degree at universities of applied sciences* or *master degree at universities of applied sciences (final examination required)*) |
|  | Other academic qualification | Other academic qualification |

Note: X indicates that we have included the aggregated group (StBA) in our analysis. Source: [72].

## Appendix 5: Concordance table of StBA with Archambault classification

| **Field classification of the StBA** | **Archambault subfield** |
| --- | --- |
| Agricultural Sciences, Food and Beverages Technology | Agronomy & Agriculture |
| Agricultural Sciences, Food and Beverages Technology | Food Science |
| Agricultural Sciences, Food and Beverages Technology | Horticulture |
| Agricultural Sciences, Food and Beverages Technology | Dairy & Animal Science |
| Biology | Toxicology |
| Biology | Bioinformatics |
| Biology | Biotechnology |
| Biology | Anthropology |
| Biology | Biochemistry & Molecular Biology |
| Biology | Biophysics |
| Biology | Developmental Biology |
| Biology | Genetics & Heredity |
| Biology | Microbiology |
| Biology | Ecology |
| Biology | Entomology |
| Biology | Evolutionary Biology |
| Biology | Marine Biology & Hydrobiology |
| Biology | Ornithology |
| Biology | Plant Biology & Botany |
| Biology | Zoology |
| Chemistry | Analytical Chemistry |
| Chemistry | General Chemistry |
| Chemistry | Inorganic & Nuclear Chemistry |
| Chemistry | Medicinal & Biomolecular Chemistry |
| Chemistry | Organic Chemistry |
| Chemistry | Physical Chemistry |
| Chemistry | Polymer |
| Chemistry | Nanoscience & Nanotechnology |
| Electrical Engineering | Optoelectronics & Photonics |
| Electrical Engineering | Electrical & Electronic Engineering |
| Electrical Engineering | Networking & Telecommunications |
| Nutrition and Home Economics | Dairy & Animal Science |
| Nutrition and Home Economics | Nutrition & Dietetics |
| Forestry, Timber Management | Forestry |
| Geosciences (excluding Geography) | Geological & Geomatics Engineering |
| Geosciences (excluding Geography) | Geochemistry & Geophysics |
| Geosciences (excluding Geography) | Geology |
| Geosciences (excluding Geography) | Meteorology & Atmospheric Sciences |
| Geosciences (excluding Geography) | Oceanography |
| Geosciences (excluding Geography) | Palaeontology |
| Geosciences (excluding Geography) | Environmental Sciences |
| Mechanical Engineering/Process Engineering | Energy |
| Mechanical Engineering/Process Engineering | Biomedical Engineering |
| Mechanical Engineering/Process Engineering | Chemical Engineering |
| Mechanical Engineering/Process Engineering | Materials |
| Mechanical Engineering/Process Engineering | Environmental Engineering |
| Mechanical Engineering/Process Engineering | Industrial Engineering & Automation |
| Mechanical Engineering/Process Engineering | Mechanical Engineering & Transports |
| Mathematics | Applied Mathematics |
| Mathematics | General Mathematics |
| Mathematics | Numerical & Comp. Mathematics |
| Mathematics | Statistics & Probability |
| Physics, Astronomy | Acoustics |
| Physics, Astronomy | Applied Physics |
| Physics, Astronomy | Astronomy & Astrophysics |
| Physics, Astronomy | Chemical Physics |
| Physics, Astronomy | Fluids & Plasmas |
| Physics, Astronomy | General Physics |
| Physics, Astronomy | Mathematical Physics |
| Physics, Astronomy | Nuclear & Particles Physics |
| Physics, Astronomy | Optics |
| Physics, Astronomy | Nanoscience & Nanotechnology |
| Psychology | Behav. Science & Comp. Psychology |
| Psychology | Clinical Psychology |
| Psychology | Developmental & Child Psychology |
| Psychology | Experimental Psychology |
| Psychology | Gene Psychology & Cognitive Sciences |
| Psychology | Human Factors |
| Psychology | Social Psychology |
| Economics | Operations Research |
| Economics | Accounting |
| Economics | Agricultural Economics & Policy |
| Economics | Business & Management |
| Economics | Development Studies |
| Economics | Econometrics |
| Economics | Economic Theory |
| Economics | Economics |
| Economics | Finance |
| Economics | Industrial Relations |
| Economics | Logistics & Transportation |
| Economics | Marketing |
| Economics | Sports, Leisure & Tourism |

## Appendix 6. Assignment of Excellence Initiative funding to subject fields

| University | Funding line | Subject field |
| --- | --- | --- |
| Albert-Ludwigs-Universität Freiburg | EC1 | Biology |
| Albert-Ludwigs-Universität Freiburg | EC2 | Biology |
| Albert-Ludwigs-Universität Freiburg | EC2 | Electrical Engineering |
| Albert-Ludwigs-Universität Freiburg | EC2 | Computer Science |
| Albert-Ludwigs-Universität Freiburg | GS1 | Biology |
| Albert-Ludwigs-Universität Freiburg | GS1 | Chemistry |
| Albert-Ludwigs-Universität Freiburg | GS1 | Medicine |
| Albert-Ludwigs-Universität Freiburg | GS2 | Biology |
| Albert-Ludwigs-Universität Freiburg | GS2 | Chemistry |
| Albert-Ludwigs-Universität Freiburg | GS2 | Medicine |
| Albert-Ludwigs-Universität Freiburg | IS1 |  |
| Carl von Ossietzky Universität Oldenburg | EC2 | Biology |
| Carl von Ossietzky Universität Oldenburg | EC2 | Medicine |
| Carl von Ossietzky Universität Oldenburg | EC2 | Physics |
| Christian-Albrechts-Universität zu Kiel | EC1 | Biology |
| Christian-Albrechts-Universität zu Kiel | EC1 | Geosciences |
| Christian-Albrechts-Universität zu Kiel | EC1 | Medicine |
| Christian-Albrechts-Universität zu Kiel | EC1 | Physics |
| Christian-Albrechts-Universität zu Kiel | EC1 | Economics |
| Christian-Albrechts-Universität zu Kiel | EC2 | Biology |
| Christian-Albrechts-Universität zu Kiel | EC2 | Geosciences |
| Christian-Albrechts-Universität zu Kiel | EC2 | Medicine |
| Christian-Albrechts-Universität zu Kiel | EC2 | Physics |
| Christian-Albrechts-Universität zu Kiel | EC2 | Economics |
| Christian-Albrechts-Universität zu Kiel | GS1 | Biology |
| Christian-Albrechts-Universität zu Kiel | GS1 | History |
| Christian-Albrechts-Universität zu Kiel | GS2 | Biology |
| Christian-Albrechts-Universität zu Kiel | GS2 | History |
| Eberhard Karls Universität Tübingen | EC1 | Medicine |
| Eberhard Karls Universität Tübingen | EC1 | Computer Science |
| Eberhard Karls Universität Tübingen | EC2 | Medicine |
| Eberhard Karls Universität Tübingen | EC2 | Computer Science |
| Eberhard Karls Universität Tübingen | GS2 | Biology |
| Eberhard Karls Universität Tübingen | GS2 | Medicine |
| Eberhard Karls Universität Tübingen | GS2 | Psychology |
| Eberhard Karls Universität Tübingen | IS2 |  |
| Freie Universität Berlin | EC1 | Classical Philology |
| Freie Universität Berlin | EC1 | History |
| Freie Universität Berlin | EC1 | Medicine |
| Freie Universität Berlin | EC1 | Psychology |
| Freie Universität Berlin | EC1 | Social Sciences |
| Freie Universität Berlin | EC2 | Classical Philology |
| Freie Universität Berlin | EC2 | History |
| Freie Universität Berlin | EC2 | Medicine |
| Freie Universität Berlin | GS1 | General and Comparative Literature Studies |
| Freie Universität Berlin | GS1 | English and American Language and Literature Studies |
| Freie Universität Berlin | GS1 | History |
| Freie Universität Berlin | GS1 | Other Language and Cultural Studies |
| Freie Universität Berlin | GS2 | General and Comparative Literature Studies |
| Freie Universität Berlin | GS2 | English and American Language and Literature Studies |
| Freie Universität Berlin | GS2 | Biology |
| Freie Universität Berlin | GS2 | Chemistry |
| Freie Universität Berlin | GS2 | History |
| Freie Universität Berlin | GS2 | Computer Science |
| Freie Universität Berlin | GS2 | Mathematics |
| Freie Universität Berlin | GS2 | Physics |
| Freie Universität Berlin | GS2 | Other Language and Cultural Studies |
| Freie Universität Berlin | IS1 |  |
| Freie Universität Berlin | IS2 |  |
| Friedrich-Alexander-Universität Erlangen-Nürnberg | EC1 | Mechanical Engineering |
| Friedrich-Alexander-Universität Erlangen-Nürnberg | EC2 | Mechanical Engineering |
| Friedrich-Alexander-Universität Erlangen-Nürnberg | GS1 | Biology |
| Friedrich-Alexander-Universität Erlangen-Nürnberg | GS1 | Chemistry |
| Friedrich-Alexander-Universität Erlangen-Nürnberg | GS1 | Physics |
| Friedrich-Alexander-Universität Erlangen-Nürnberg | GS2 | Biology |
| Friedrich-Alexander-Universität Erlangen-Nürnberg | GS2 | Chemistry |
| Friedrich-Alexander-Universität Erlangen-Nürnberg | GS2 | Physics |
| Friedrich-Schiller-Universität Jena | GS1 | Biology |
| Friedrich-Schiller-Universität Jena | GS2 | Biology |
| Georg-August-Universität Göttingen | EC1 | Biology |
| Georg-August-Universität Göttingen | EC1 | Chemistry |
| Georg-August-Universität Göttingen | EC1 | Physics |
| Georg-August-Universität Göttingen | EC2 | Biology |
| Georg-August-Universität Göttingen | EC2 | Chemistry |
| Georg-August-Universität Göttingen | EC2 | Physics |
| Georg-August-Universität Göttingen | GS1 | Biology |
| Georg-August-Universität Göttingen | GS1 | Chemistry |
| Georg-August-Universität Göttingen | GS2 | Biology |
| Georg-August-Universität Göttingen | GS2 | Chemistry |
| Georg-August-Universität Göttingen | IS1 |  |
| Goethe-Universität Frankfurt am Main | EC1 | Biology |
| Goethe-Universität Frankfurt am Main | EC1 | Chemistry |
| Goethe-Universität Frankfurt am Main | EC1 | History |
| Goethe-Universität Frankfurt am Main | EC1 | Medicine |
| Goethe-Universität Frankfurt am Main | EC1 | Philosophy |
| Goethe-Universität Frankfurt am Main | EC1 | Political Science |
| Goethe-Universität Frankfurt am Main | EC2 | Biology |
| Goethe-Universität Frankfurt am Main | EC2 | Chemistry |
| Goethe-Universität Frankfurt am Main | EC2 | History |
| Goethe-Universität Frankfurt am Main | EC2 | Medicine |
| Goethe-Universität Frankfurt am Main | EC2 | Philosophy |
| Goethe-Universität Frankfurt am Main | EC2 | Political Science |
| Gottfried Wilhelm Leibniz Universität Hannover | EC1 | Physics |
| Gottfried Wilhelm Leibniz Universität Hannover | EC2 | Biology |
| Gottfried Wilhelm Leibniz Universität Hannover | EC2 | Medicine |
| Gottfried Wilhelm Leibniz Universität Hannover | EC2 | Physics |
| Heinrich-Heine-Universität Düsseldorf | EC2 | Biology |
| Humboldt-Universität zu Berlin | EC1 | Classical Philology |
| Humboldt-Universität zu Berlin | EC1 | History |
| Humboldt-Universität zu Berlin | EC1 | Medicine |
| Humboldt-Universität zu Berlin | EC2 | Classical Philology |
| Humboldt-Universität zu Berlin | EC2 | History |
| Humboldt-Universität zu Berlin | EC2 | Medicine |
| Humboldt-Universität zu Berlin | GS1 | General and Comparative Literature Studies |
| Humboldt-Universität zu Berlin | GS1 | Biology |
| Humboldt-Universität zu Berlin | GS1 | Chemistry |
| Humboldt-Universität zu Berlin | GS1 | Philosophy |
| Humboldt-Universität zu Berlin | GS1 | Physics |
| Humboldt-Universität zu Berlin | GS1 | Political Science |
| Humboldt-Universität zu Berlin | GS1 | Psychology |
| Humboldt-Universität zu Berlin | GS1 | Social Sciences |
| Humboldt-Universität zu Berlin | GS2 | General and Comparative Literature Studies |
| Humboldt-Universität zu Berlin | GS2 | Biology |
| Humboldt-Universität zu Berlin | GS2 | Chemistry |
| Humboldt-Universität zu Berlin | GS2 | History |
| Humboldt-Universität zu Berlin | GS2 | Computer Science |
| Humboldt-Universität zu Berlin | GS2 | Mathematics |
| Humboldt-Universität zu Berlin | GS2 | Philosophy |
| Humboldt-Universität zu Berlin | GS2 | Physics |
| Humboldt-Universität zu Berlin | GS2 | Psychology |
| Humboldt-Universität zu Berlin | GS2 | Other Language and Cultural Studies |
| Humboldt-Universität zu Berlin | IS2 |  |
| Johannes Gutenberg-Universität Mainz | EC2 | Physics |
| Johannes Gutenberg-Universität Mainz | GS1 | Physics |
| Johannes Gutenberg-Universität Mainz | GS2 | Physics |
| Julius-Maximilians-Universität Würzburg | GS1 | Biology |
| Julius-Maximilians-Universität Würzburg | GS1 | Chemistry |
| Julius-Maximilians-Universität Würzburg | GS1 | Medicine |
| Julius-Maximilians-Universität Würzburg | GS2 | Biology |
| Julius-Maximilians-Universität Würzburg | GS2 | Chemistry |
| Julius-Maximilians-Universität Würzburg | GS2 | Medicine |
| Justus-Liebig-Universität Gießen | EC1 | Medicine |
| Justus-Liebig-Universität Gießen | EC2 | Medicine |
| Justus-Liebig-Universität Gießen | GS1 | English and American Language and Literature Studies |
| Justus-Liebig-Universität Gießen | GS1 | Language and Cultural Studies, general |
| Justus-Liebig-Universität Gießen | GS2 | Anglistik und Amerikanistik |
| Justus-Liebig-Universität Gießen | GS2 | English and American Language and Literature Studies |
| Karlsruher Institut für Technologie | EC1 | Physics |
| Karlsruher Institut für Technologie | GS1 | Physics |
| Karlsruher Institut für Technologie | GS2 | Physics |
| Karlsruher Institut für Technologie | IS1 |  |
| Ludwig-Maximilians-Universität München | EC1 | Biology |
| Ludwig-Maximilians-Universität München | EC1 | Chemistry |
| Ludwig-Maximilians-Universität München | EC1 | Physics |
| Ludwig-Maximilians-Universität München | EC2 | Biology |
| Ludwig-Maximilians-Universität München | EC2 | Chemistry |
| Ludwig-Maximilians-Universität München | EC2 | Medicine |
| Ludwig-Maximilians-Universität München | EC2 | Physics |
| Ludwig-Maximilians-Universität München | GS1 | Biology |
| Ludwig-Maximilians-Universität München | GS2 | Classical Philology |
| Ludwig-Maximilians-Universität München | GS2 | Biology |
| Ludwig-Maximilians-Universität München | GS2 | Medicine |
| Ludwig-Maximilians-Universität München | GS2 | Other Language and Cultural Studies |
| Ludwig-Maximilians-Universität München | IS1 |  |
| Ludwig-Maximilians-Universität München | IS2 |  |
| Otto-Friedrich-Universität Bamberg | GS2 | Political Science |
| Otto-Friedrich-Universität Bamberg | GS2 | Social Sciences |
| Rheinische Friedrich-Wilhelms-Universität Bonn | EC1 | Mathematics |
| Rheinische Friedrich-Wilhelms-Universität Bonn | EC2 | Biology |
| Rheinische Friedrich-Wilhelms-Universität Bonn | EC2 | Mathematics |
| Rheinische Friedrich-Wilhelms-Universität Bonn | GS1 | Physics |
| Rheinische Friedrich-Wilhelms-Universität Bonn | GS1 | Economics |
| Rheinische Friedrich-Wilhelms-Universität Bonn | GS2 | Physics |
| Rheinisch-Westfälische Technische Hochschule Aachen | EC1 | Electrical Engineering |
| Rheinisch-Westfälische Technische Hochschule Aachen | EC1 | Mechanical Engineering |
| Rheinisch-Westfälische Technische Hochschule Aachen | EC2 | Mechanical Engineering |
| Rheinisch-Westfälische Technische Hochschule Aachen | GS1 | Computer Science |
| Rheinisch-Westfälische Technische Hochschule Aachen | GS2 | Computer Science |
| Rheinisch-Westfälische Technische Hochschule Aachen | IS1 |  |
| Rheinisch-Westfälische Technische Hochschule Aachen | IS2 |  |
| Ruhr-Universität Bochum | EC2 | Chemistry |
| Ruprecht-Karls-Universität Heidelberg | EC1 | Biology |
| Ruprecht-Karls-Universität Heidelberg | EC1 | Chemistry |
| Ruprecht-Karls-Universität Heidelberg | EC1 | Cultural Studies in the narrow sense |
| Ruprecht-Karls-Universität Heidelberg | EC1 | Other Language and Cultural Studies |
| Ruprecht-Karls-Universität Heidelberg | EC2 | Medicine |
| Ruprecht-Karls-Universität Heidelberg | EC2 | Cultural Studies in the narrow sense |
| Ruprecht-Karls-Universität Heidelberg | EC2 | Other Language and Cultural Studies |
| Ruprecht-Karls-Universität Heidelberg | GS1 | Biology |
| Ruprecht-Karls-Universität Heidelberg | GS1 | Medicine |
| Ruprecht-Karls-Universität Heidelberg | GS1 | Mathematics |
| Ruprecht-Karls-Universität Heidelberg | GS1 | Physics |
| Ruprecht-Karls-Universität Heidelberg | GS2 | Biology |
| Ruprecht-Karls-Universität Heidelberg | GS2 | Medicine |
| Ruprecht-Karls-Universität Heidelberg | GS2 | Mathematics |
| Ruprecht-Karls-Universität Heidelberg | GS2 | Physics |
| Ruprecht-Karls-Universität Heidelberg | IS1 |  |
| Ruprecht-Karls-Universität Heidelberg | IS2 |  |
| Technische Universität Berlin | EC1 | Chemistry |
| Technische Universität Berlin | EC2 | Chemistry |
| Technische Universität Berlin | GS1 | Mathematics |
| Technische Universität Berlin | GS2 | Mathematics |
| Technische Universität Chemnitz | EC2 | Mechanical Engineering |
| Technische Universität Darmstadt | EC1 | Mechanical Engineering |
| Technische Universität Darmstadt | GS1 | Computer Science |
| Technische Universität Darmstadt | GS1 | Mathematics |
| Technische Universität Darmstadt | GS2 | Computer Science |
| Technische Universität Darmstadt | GS2 | Mechanical Engineering |
| Technische Universität Darmstadt | GS2 | Mathematics |
| Technische Universität Dresden | EC1 | Biology |
| Technische Universität Dresden | EC2 | Biology |
| Technische Universität Dresden | EC2 | Electrical Engineering |
| Technische Universität Dresden | EC2 | Computer Science |
| Technische Universität Dresden | GS1 | Biology |
| Technische Universität Dresden | GS1 | Computer Science |
| Technische Universität Dresden | GS1 | Mechanical Engineering |
| Technische Universität Dresden | GS2 | Biology |
| Technische Universität Dresden | GS2 | Computer Science |
| Technische Universität Dresden | GS2 | Mechanical Engineering |
| Technische Universität Dresden | IS2 |  |
| Technische Universität Kaiserslautern | GS2 | Physics |
| Technische Universität München | EC1 | Biology |
| Technische Universität München | EC1 | Chemistry |
| Technische Universität München | EC1 | Mechanical Engineering |
| Technische Universität München | EC1 | Physics |
| Technische Universität München | EC2 | Biology |
| Technische Universität München | EC2 | Chemistry |
| Technische Universität München | EC2 | Medicine |
| Technische Universität München | EC2 | Physics |
| Technische Universität München | IS1 |  |
| Technische Universität München | IS2 |  |
| Universität Augsburg | EC2 | Physics |
| Universität Bayreuth | GS1 | Other Language and Cultural Studies |
| Universität Bayreuth | GS2 | Other Language and Cultural Studies |
| Universität Bielefeld | EC1 | Biology |
| Universität Bielefeld | EC1 | Computer Science |
| Universität Bielefeld | EC1 | Sports Science |
| Universität Bielefeld | EC2 | Biology |
| Universität Bielefeld | EC2 | Computer Science |
| Universität Bielefeld | EC2 | Sports Science |
| Universität Bielefeld | GS1 | History |
| Universität Bielefeld | GS1 | Philosophy |
| Universität Bielefeld | GS1 | Social Sciences |
| Universität Bielefeld | GS2 | History |
| Universität Bielefeld | GS2 | Philosophy |
| Universität Bielefeld | GS2 | Social Sciences |
| Universität Bremen | EC1 | Geosciences |
| Universität Bremen | EC2 | Geosciences |
| Universität Bremen | GS1 | Biology |
| Universität Bremen | GS1 | Chemistry |
| Universität Bremen | GS1 | Political Science |
| Universität Bremen | GS1 | Psychology |
| Universität Bremen | GS1 | Social Sciences |
| Universität Bremen | GS2 | Political Science |
| Universität Bremen | GS2 | Psychology |
| Universität Bremen | GS2 | Social Sciences |
| Universität Bremen | IS2 |  |
| Universität des Saarlandes | EC1 | Computer Science |
| Universität des Saarlandes | EC2 | Computer Science |
| Universität des Saarlandes | GS1 | Computer Science |
| Universität des Saarlandes | GS2 | Computer Science |
| Universität Hamburg | EC1 | Geosciences |
| Universität Hamburg | EC2 | Chemistry |
| Universität Hamburg | EC2 | Geosciences |
| Universität Hamburg | EC2 | Physics |
| Universität Konstanz | EC1 | Philosophy |
| Universität Konstanz | EC1 | Political Science |
| Universität Konstanz | EC1 | Sociology |
| Universität Konstanz | EC2 | Philosophy |
| Universität Konstanz | EC2 | Political Science |
| Universität Konstanz | EC2 | Sociology |
| Universität Konstanz | GS1 | Biology |
| Universität Konstanz | GS1 | Chemistry |
| Universität Konstanz | GS2 | Biology |
| Universität Konstanz | GS2 | Chemistry |
| Universität Konstanz | GS2 | Political Science |
| Universität Konstanz | GS2 | Psychology |
| Universität Konstanz | GS2 | Economics |
| Universität Konstanz | IS1 |  |
| Universität Konstanz | IS2 |  |
| Universität Leipzig | GS1 | Physics |
| Universität Mannheim | GS1 | Social Sciences |
| Universität Mannheim | GS1 | Economics |
| Universität Mannheim | GS2 | Social Sciences |
| Universität Mannheim | GS2 | Economics |
| Universität Regensburg | GS2 | Other Language and Cultural Studies |
| Universität Stuttgart | GS1 | Computer Science |
| Universität Stuttgart | GS1 | Economics |
| Universität Stuttgart | GS2 | Computer Science |
| Universität Stuttgart | GS2 | Economics |
| Universität Ulm | GS1 | Medicine |
| Universität Ulm | GS2 | Medicine |
| Universität zu Köln | EC1 | Biology |
| Universität zu Köln | EC2 | Biology |
| Universität zu Köln | GS1 | Physics |
| Universität zu Köln | GS2 | Philosophy |
| Universität zu Köln | GS2 | Physics |
| Universität zu Köln | IS2 |  |
| Westfälische Wilhelms-Universität Münster | EC1 | Protestant Theology |
| Westfälische Wilhelms-Universität Münster | EC1 | Catholic Theology |
| Westfälische Wilhelms-Universität Münster | EC1 | Political Science |
| Westfälische Wilhelms-Universität Münster | EC2 | Biology |
| Westfälische Wilhelms-Universität Münster | EC2 | Chemistry |
| Westfälische Wilhelms-Universität Münster | EC2 | Protestant Theology |
| Westfälische Wilhelms-Universität Münster | EC2 | Medicine |
| Westfälische Wilhelms-Universität Münster | EC2 | Catholic Theology |
| Westfälische Wilhelms-Universität Münster | EC2 | Political Science |

Note: ECx are clusters of excellence; GSx are Graduate Research Schools; ISx are institutional strategies. The index x refers to the funding phase; x=1 is the first funding phase 2006-2011, x=2 the second phase 2012- 2017. The field “medicine” is for information only, because we did not process data on medical faculties due to the data problems mentioned in Section 3.

## Appendix 7: Correlation matrices

**Tab. 7a:** Correlation matrix, first “initiative” phase (2006-2011), all universities.

|  | Professors | Total grant funding | DFG grant funding | Students | Citations |
| --- | --- | --- | --- | --- | --- |
| Professors | 1.000 |  |  |  |  |
| Total grant funding | 0.578*** | 1.000 |  |  |  |
| DFG grant funding | 0.203*** | 0.215*** | 1.000 |  |  |
| Students | 0.688*** | 0.352*** | 0.201*** | 1.000 |  |
| Citations | 0.436*** | 0.438*** | 0.159*** | 0.106*** | 1.000 |
| Observations | 2,388 |  |  |  |  |

\* $p < 0.05$ , \*\* $p < 0.01$ , \*\*\* $p < 0.001$

**Tab. 7b:** Correlation matrix, second “initiative” phase (2012-2017), all universities.

|  | Professors | Total grant funding | DFG grant funding | Students | Citations | Phase 1 |
| --- | --- | --- | --- | --- | --- | --- |
| Professors | 1.000 |  |  |  |  |  |
| Total grant funding | 0.607*** | 1.000 |  |  |  |  |
| DFG grant funding | 0.232*** | 0.214*** | 1.000 |  |  |  |
| Students | 0.730*** | 0.492*** | 0.195*** | 1.000 |  |  |
| Citations | 0.477*** | 0.477*** | 0.176*** | 0.159*** | 1.000 |  |
| Phase 1 | 0.329*** | 0.365*** | 0.179*** | 0.180*** | 0.430*** | 1.000 |
| Observations | 2,396 |  |  |  |  |  |

\* $p < 0.05$ , \*\* $p < 0.01$ , \*\*\* $p < 0.001$

**Tab. 7c:**
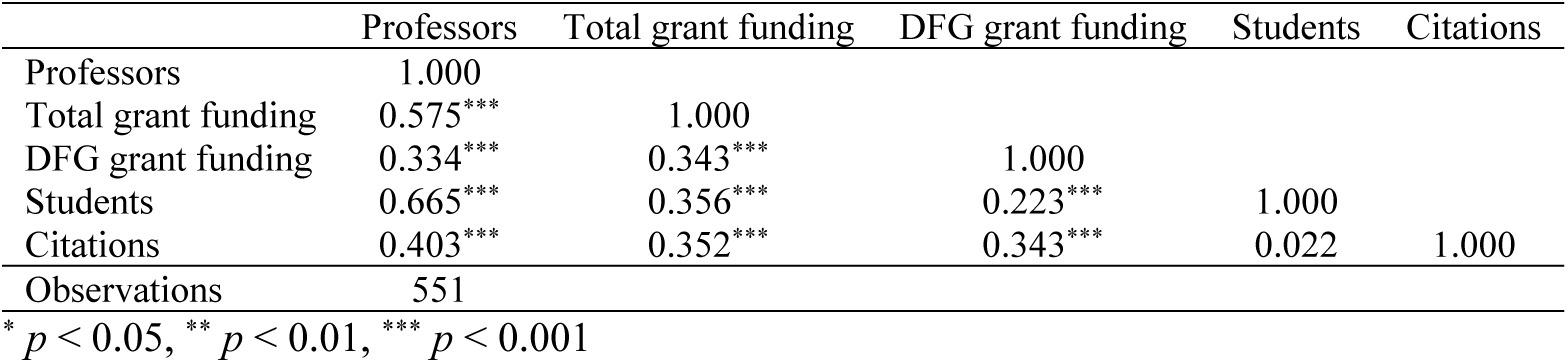
Correlation matrix, first “initiative” phase (2006-2011), 12 subject fields with good WoS coverage, all universities.

**Tab. 7d:**
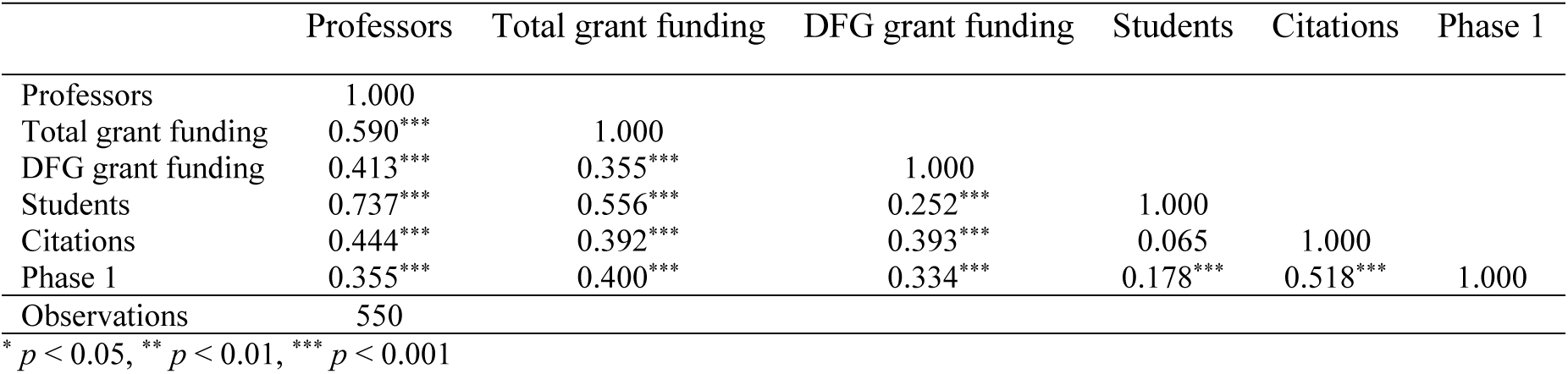
Correlation matrix, second “initiative” phase (2012-2017), 12 subject fields with good WoS coverage, all universities.

|  | Professors | Total grant funding | DFG grant funding | Students | Citations | Phase 1 |
| --- | --- | --- | --- | --- | --- | --- |
| Professors | 1.000 |  |  |  |  |  |
| Total grant funding | 0.590*** | 1.000 |  |  |  |  |
| DFG grant funding | 0.413*** | 0.355*** | 1.000 |  |  |  |
| Students | 0.737*** | 0.556*** | 0.252*** | 1.000 |  |  |
| Citations | 0.444*** | 0.392*** | 0.393*** | 0.065 | 1.000 |  |
| Phase 1 | 0.355*** | 0.400*** | 0.334*** | 0.178*** | 0.518*** | 1.000 |
| Observations | 550 |  |  |  |  |  |

\* $p < 0.05$ , \*\* $p < 0.01$ , \*\*\* $p < 0.001$

## Appendix 8: Logistic regression analyses

**Tab. 8a:**
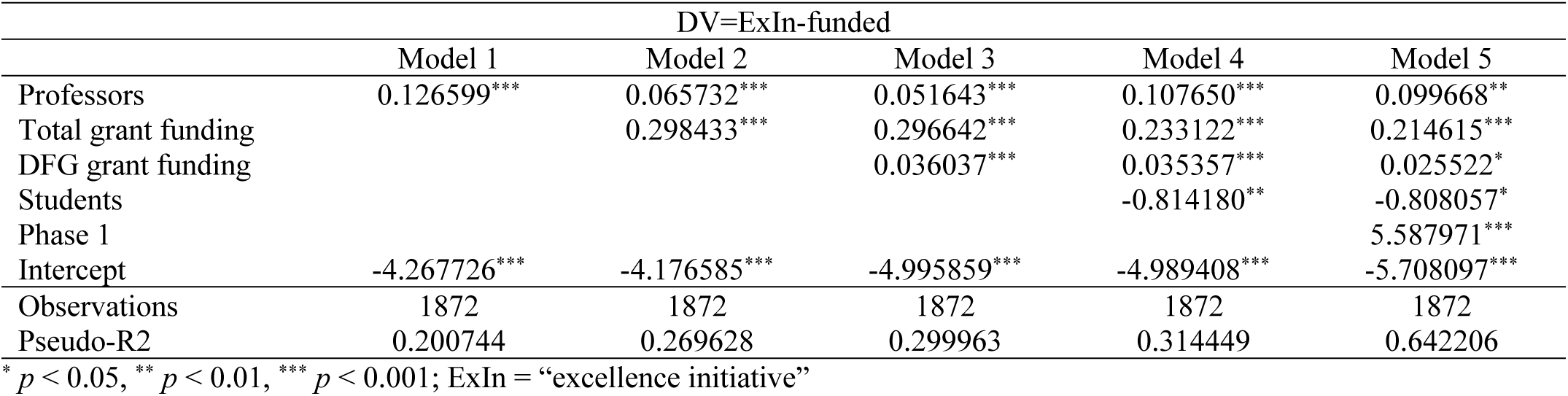
Logistic regression, second “initiative” phase (2012-2017), NTUs.

|  | DV=ExIn-funded |  |  |  |  |
| --- | --- | --- | --- | --- | --- |
|  | Model 1 | Model 2 | Model 3 | Model 4 | Model 5 |
| Professors | 0.126599*** | 0.065732*** | 0.051643*** | 0.107650*** | 0.099668** |
| Total grant funding |  | 0.298433*** | 0.296642*** | 0.233122*** | 0.214615*** |
| DFG grant funding |  |  | 0.036037*** | 0.035357*** | 0.025522* |
| Students |  |  |  | -0.814180** | -0.808057* |
| Phase 1 |  |  |  |  | 5.587971*** |
| Intercept | -4.267726*** | -4.176585*** | -4.995859*** | -4.989408*** | -5.708097*** |
| Observations | 1872 | 1872 | 1872 | 1872 | 1872 |
| Pseudo-R2 | 0.200744 | 0.269628 | 0.299963 | 0.314449 | 0.642206 |

\* $p < 0.05$ , \*\* $p < 0.01$ , \*\*\* $p < 0.001$ ; ExIn = “excellence initiative”

**Tab. 8b:** Logistic regression, second “initiative” phase (2012-2017), TUs.

|  | DV=ExIn-funded |  |  |  |  |
| --- | --- | --- | --- | --- | --- |
|  | Model 1 | Model 2 | Model 3 | Model 4 | Model 5 |
| Professors | 0.145839*** | 0.129660*** | 0.126906*** | 0.170633*** | 0.210778** |
| Total grant funding |  | 0.022429 | 0.021843 | 0.105880* | 0.062580 |
| DFG grant funding |  |  | 0.006038 | 0.007909 | -0.059053 |
| Students |  |  |  | -1.332431* | -1.755588 |
| Phase 1 |  |  |  |  | 7.673918*** |
| Intercept | -5.314181*** | -5.194264*** | -5.356676*** | -5.548694*** | -5.340610*** |
| Observations | 524 | 524 | 524 | 524 | 524 |
| Pseudo-R2 | 0.318055 | 0.321262 | 0.322242 | 0.359837 | 0.746805 |

\* $p < 0.05$ , \*\* $p < 0.01$ , \*\*\* $p < 0.001$ ; ExIn = “excellence initiative”

**Tab. 8c:** Logistic regression, first “initiative” phase (2006-2011), NTUs.

|  | DV=ExIn-funded |  |  |  |
| --- | --- | --- | --- | --- |
|  | Model 1 | Model 2 | Model 3 | Model 4 |
| Professors | 0.102579*** | 0.055361*** | 0.047246*** | 0.060696** |
| Total grant funding |  | 0.350526*** | 0.313880*** | 0.289041*** |
| DFG grant funding |  |  | 0.081932*** | 0.083187*** |
| Students |  |  |  | -0.175910 |
| Intercept | -4.145971*** | -4.154941*** | -5.352402*** | -5.367308*** |
| Observations | 1852 | 1852 | 1852 | 1852 |
| Pseudo-R2 | 0.132634 | 0.194125 | 0.236820 | 0.238270 |

\* $p < 0.05$ , \*\* $p < 0.01$ , \*\*\* $p < 0.001$ ; ExIn = “excellence initiative”

**Tab. 8d:** Logistic regression, first “initiative” phase (2006-2011), TUs.

|  | DV=ExIn-funded |  |  |  |
| --- | --- | --- | --- | --- |
|  | Model 1 | Model 2 | Model 3 | Model 4 |
| Professors | 0.121399*** | 0.098810*** | 0.101597*** | 0.085094** |
| Total grant funding |  | 0.051689 | 0.035792 | -0.007922 |
| DFG grant funding |  |  | 0.060482 | 0.054749 |
| Students |  |  |  | 0.869810* |
| Intercept | -4.964031*** | -4.837065*** | -6.132478*** | -6.400819*** |
| Observations | 536 | 536 | 536 | 536 |
| Pseudo-R2 | 0.254066 | 0.264581 | 0.288912 | 0.319044 |

\* $p < 0.05$ , \*\* $p < 0.01$ , \*\*\* $p < 0.001$ ; ExIn = “excellence initiative”

**Tab. 8e:** Logistic regression, first “initiative” phase (2006-2011), natural sciences.

|  | DV=ExIn-funded |  |  |  |
| --- | --- | --- | --- | --- |
|  | Model 1 | Model 2 | Model 3 | Model 4 |
| Professors | 0.155950*** | 0.082983** | 0.077637** | 0.075690** |
| Total grant funding |  | 0.375925*** | 0.321290*** | 0.320879*** |
| DFG grant funding |  |  | 0.054876** | 0.054596* |
| Students |  |  |  | 0.071758 |
| Intercept | -4.543776*** | -4.534415*** | -5.285426*** | -5.295774*** |
| Observations | 514 | 514 | 514 | 514 |
| Pseudo-R2 | 0.235079 | 0.309631 | 0.328897 | 0.329021 |

\* $p < 0.05$ , \*\* $p < 0.01$ , \*\*\* $p < 0.001$ ; ExIn = “excellence initiative”

**Tab. 8f:** Logistic regression, second “initiative” phase (2012-2017), natural sciencesTab. 8g: Logistic regression, first “initiative” phase (2006-2011), social sciences.

|  | DV=ExIn-funded |  |  |  |  |
| --- | --- | --- | --- | --- | --- |
|  | Model 1 | Model 2 | Model 3 | Model 4 | Model 5 |
| Professors | 0.215851*** | 0.142938*** | 0.134247*** | 0.147732*** | 0.184459*** |
| Total grant funding |  | 0.221826*** | 0.213142*** | 0.210432*** | 0.088022 |
| DFG grant funding |  |  | 0.012736 | 0.013054 | -0.002499 |
| Students |  |  |  | -0.377013 | -0.299023 |
| Phase 1 |  |  |  |  | 4.729829*** |
| Intercept | -5.421236*** | -5.111836*** | -5.295268*** | -5.259958*** | -6.038445*** |
| Observations | 512 | 512 | 512 | 512 | 512 |
| Pseudo-R2 | 0.374617 | 0.415087 | 0.418316 | 0.420407 | 0.651497 |

\* $p < 0.05$ , \*\* $p < 0.01$ , \*\*\* $p < 0.001$ ; ExIn = “excellence initiative”

**Tab. 8g:** Logistic regression, first “initiative” phase (2006-2011), social sciences.

|  | DV=ExIn-funded |  |  |  |
| --- | --- | --- | --- | --- |
|  | Model 1 | Model 2 | Model 3 | Model 4 |
| Professors | 0.052552** | 0.034296 | 0.035050 | 0.090940 |
| Total grant funding |  | 0.430796* | 0.363529 | 0.378447* |
| DFG grant funding |  |  | 0.031375 | 0.036132 |
| Students |  |  |  | -0.537127 |
| Intercept | -3.826455*** | -3.933624*** | -4.389794*** | -4.493444*** |
| Observations | 431 | 431 | 431 | 431 |
| Pseudo-R2 | 0.043616 | 0.073950 | 0.082885 | 0.095758 |

\* $p < 0.05$ , \*\* $p < 0.01$ , \*\*\* $p < 0.001$ ; ExIn = “excellence initiative”

**Tab. 8h:** Logistic regression, second “initiative” phase (2012-2017), social sciences.

|  | DV=ExIn-funded |  |  |  |  |
| --- | --- | --- | --- | --- | --- |
|  | Model 1 | Model 2 | Model 3 | Model 4 | Model 5 |
| Professors | 0.049424** | 0.018250 | 0.018541 | 0.129890* | 0.053212 |
| Total grant funding |  | 0.383354 | 0.397720 | 0.543396* | -0.102404 |
| DFG grant funding |  |  | -0.005972 | -0.008393 | -0.096110 |
| Students |  |  |  | -1.643999* | -0.154471 |
| Phase 1 |  |  |  |  | 7.680742*** |
| Intercept | -3.913044*** | -3.906463*** | -3.781269*** | -3.676088*** | -3.878507*** |
| Observations | 435 | 435 | 435 | 435 | 435 |
| Pseudo-R2 | 0.045639 | 0.067744 | 0.068850 | 0.119069 | 0.617039 |

\* $p < 0.05$ , \*\* $p < 0.01$ , \*\*\* $p < 0.001$ ; ExIn = “excellence initiative”

**Tab. 8i:** Logistic regression, first “initiative” phase (2006-2011), humanities.

|  | DV=ExIn-funded |  |  |  |
| --- | --- | --- | --- | --- |
|  | Model 1 | Model 2 | Model 3 | Model 4 |
| Professors | 0.096919*** | 0.066854* | 0.059251 | 0.058412 |
| Total grant funding |  | 0.593877* | 0.502999 | 0.501938 |
| DFG grant funding |  |  | 0.053168* | 0.053203* |
| Students |  |  |  | 0.013085 |
| Intercept | -4.229057*** | -4.277242*** | -5.082191*** | -5.083426*** |
| Observations | 892 | 892 | 892 | 892 |
| Pseudo-R2 | 0.055673 | 0.073600 | 0.095725 | 0.095733 |

\* $p < 0.05$ , \*\* $p < 0.01$ , \*\*\* $p < 0.001$ ; ExIn = “excellence initiative”

**Tab. 8j:**
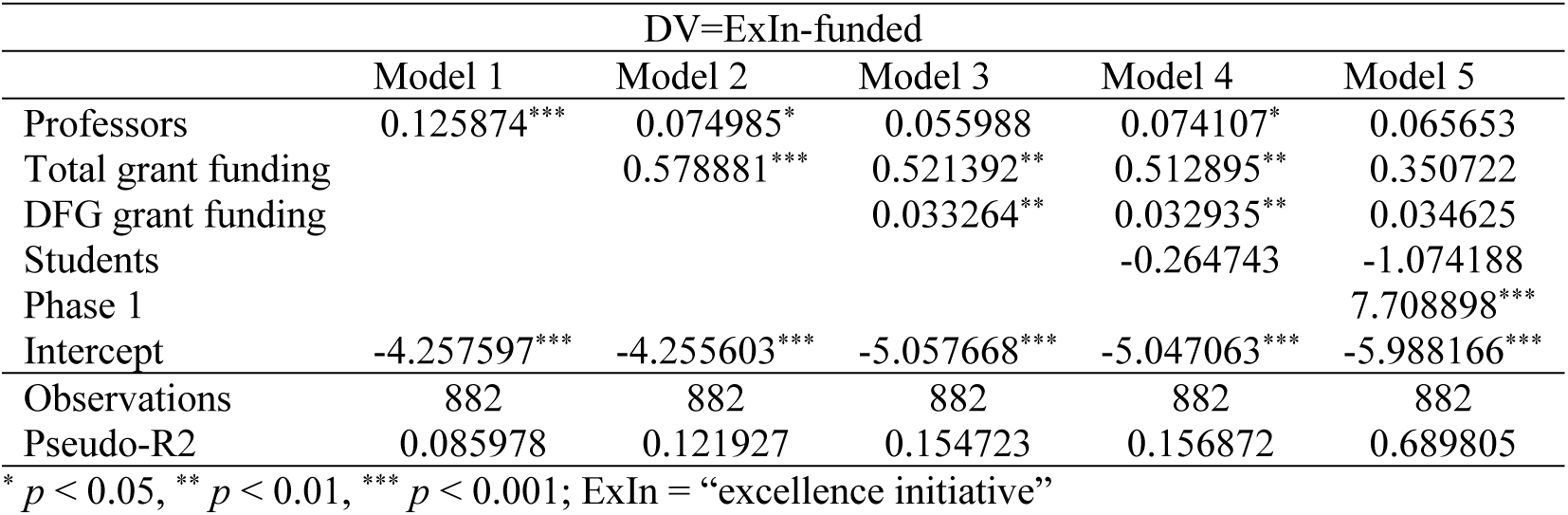
Logistic regression, second “initiative” phase (2012-2017), humanities.

**Tab. 8k:** Logistic regression, first “initiative” phase (2006-2011), 12 subject fields with good WoS coverage, all universities.

|  | DV=ExIn-funded |  |  |  |  |
| --- | --- | --- | --- | --- | --- |
|  | Model 1 | Model 2 | Model 3 | Model 4 | Model 5 |
| Citations | 0.491121*** | 0.423491*** | 0.394527*** | 0.371862*** | 0.399359*** |
| DFG grant funding |  | 0.080448*** | 0.065535*** | 0.065521*** | 0.064095** |
| Total grant funding |  |  | 0.082459* | 0.058416 | 0.058690 |
| Professors |  |  |  | 0.024477 | 0.009057 |
| Students |  |  |  |  | 0.226920 |
| Intercept | -3.120918*** | -4.442549*** | -4.432054*** | -4.732205*** | -4.680096*** |
| Observations | 551 | 551 | 551 | 551 | 551 |
| Pseudo-R2 | 0.242012 | 0.293836 | 0.314653 | 0.318042 | 0.320264 |

\* $p < 0.05$ , \*\* $p < 0.01$ , \*\*\* $p < 0.001$ ; ExIn = “excellence initiative”

**Tab. 8l:**
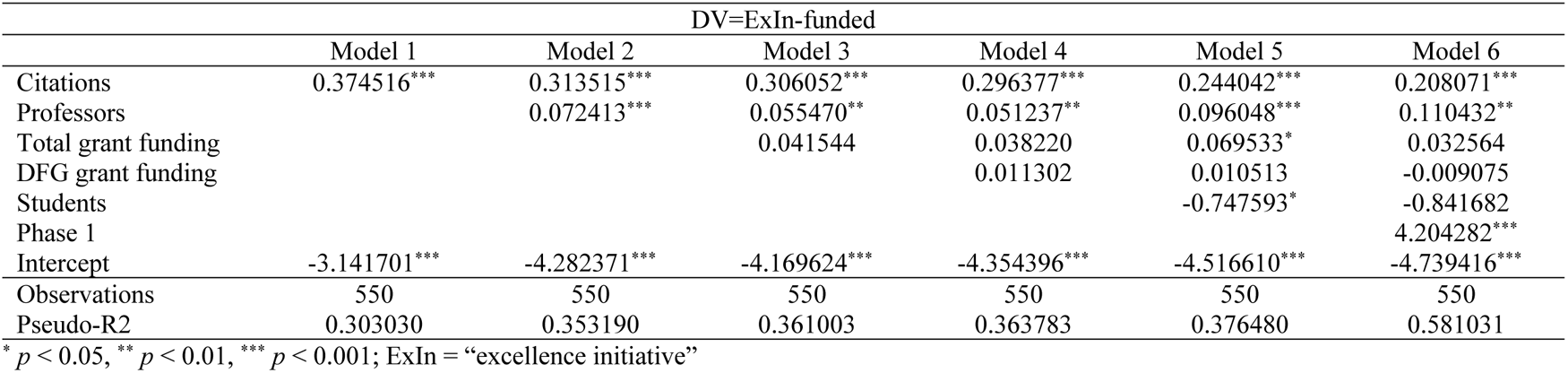
Logistic regression, second “initiative” phase (2012-2017), 12 subject fields with good WoS coverage, all universities.

## Appendix 9: Two-level logistic regression analyses

**Tab. 9a:**
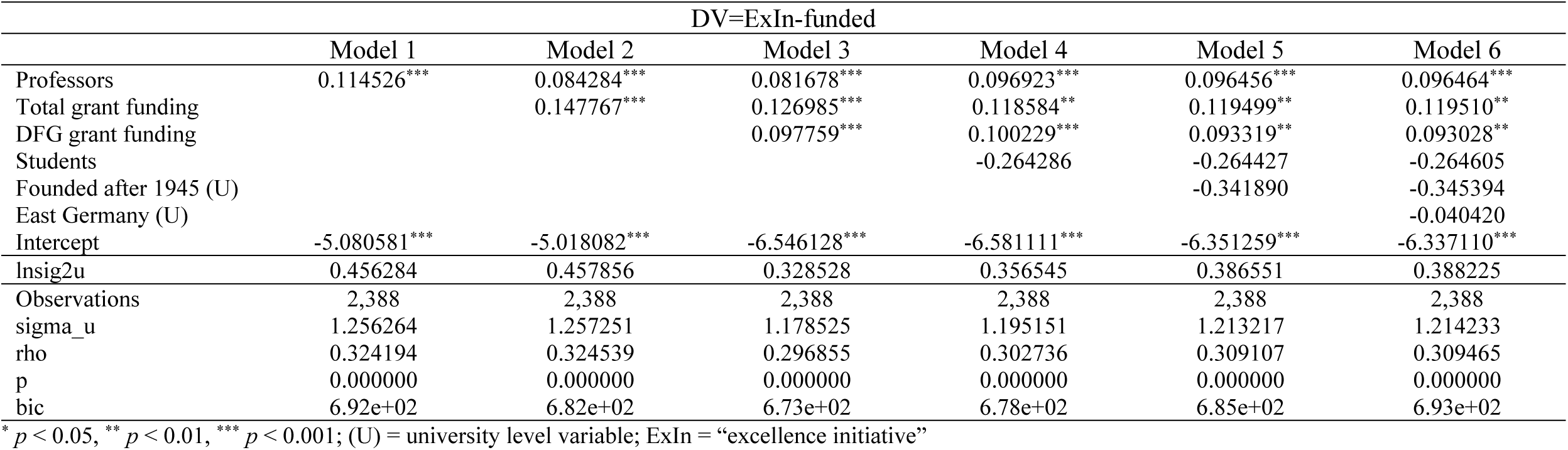
Two-level logistic regression, first “initiative” phase (2006-2011), all universities.

**Tab. 9b:** Two-level logistic regression, second “initiative” phase (2012-2017), all universities.

|  | DV=ExIn-funded |  |  |  |  |  |  |
| --- | --- | --- | --- | --- | --- | --- | --- |
|  | Model 1 | Model 2 | Model 3 | Model 4 | Model 5 | Model 6 | Model 7 |
| Professors | 0.134104*** | 0.098283*** | 0.093758*** | 0.167302*** | 0.171687*** | 0.173286*** | 0.174681*** |
| Total grant funding |  | 0.119472*** | 0.111851*** | 0.124381*** | 0.070207* | 0.068659 | 0.071790* |
| DFG grant funding |  |  | 0.031678** | 0.032247** | 0.004028 | 0.012131 | 0.011722 |
| Students |  |  |  | -1.394023*** | -1.541516*** | -1.529333*** | -1.554121*** |
| Phase 1 |  |  |  |  | 6.040811*** | 6.048217*** | 6.073146*** |
| Founded after 1945 (U) |  |  |  |  |  | 0.640197 | 0.638906 |
| East Germany (U) |  |  |  |  |  |  | -0.332986 |
| Intercept | -4.931936*** | -4.912985*** | -5.657228*** | -5.756519*** | -5.738849*** | -6.242998*** | -6.203378*** |
| lnsig2u | -0.029492 | 0.201887 | 0.108190 | 0.171891 | -0.506117 | -0.637020 | -0.506778 |
| Observations | 2,396 | 2,396 | 2,396 | 2,396 | 2,396 | 2,396 | 2,396 |
| sigma_u | 0.985362 | 1.106214 | 1.055585 | 1.089747 | 0.776422 | 0.727232 | 0.776166 |
| rho | 0.227877 | 0.271117 | 0.253003 | 0.265231 | 0.154862 | 0.138493 | 0.154776 |
| p | 0.000000 | 0.000000 | 0.000000 | 0.000000 | 0.000000 | 0.000000 | 0.000000 |
| bic | 7.60e+02 | 7.44e+02 | 7.43e+02 | 7.09e+02 | 4.04e+02 | 4.10e+02 | 4.17e+02 |

\* $p < 0.05$ , \*\* $p < 0.01$ , \*\*\* $p < 0.001$ ; (U) = university level variable; ExIn = “excellence initiative”

**Tab. 9c:** Two-level logistic regression, first “initiative” phase (2006-2011), 12 subject fields with good WoS coverage, all universities.

|  | DV=ExIn-funded |  |  |  |  |  |  |
| --- | --- | --- | --- | --- | --- | --- | --- |
|  | Model 1 | Model 2 | Model 3 | Model 4 | Model 5 | Model 6 | Model 7 |
| Citations | 0.566880*** | 0.494364*** | 0.492519*** | 0.445917*** | 0.500020*** | 0.495780*** | 0.505399*** |
| Professors |  | 0.062377** | 0.030574 | 0.026949 | 0.001290 | -0.001131 | -0.003900 |
| Total grant funding |  |  | 0.085760* | 0.066730 | 0.066973 | 0.068932 | 0.069469 |
| DFG grant funding |  |  |  | 0.090206** | 0.089567** | 0.070416* | 0.074635* |
| Students |  |  |  |  | 0.368885 | 0.372392 | 0.398133 |
| Founded after 1945 (U) |  |  |  |  |  | -0.960572 | -0.906028 |
| East Germany (U) |  |  |  |  |  |  | 0.468667 |
| Intercept | -3.802962*** | -4.767178*** | -4.516534*** | -5.837735*** | -5.826572*** | -5.154399*** | -5.343520*** |
| lnsig2u | 0.381569 | 0.380839 | 0.331702 | 0.211194 | 0.294432 | 0.282858 | 0.268067 |
| Observations | 551 | 551 | 551 | 551 | 551 | 551 | 551 |
| sigma_u | 1.210199 | 1.209757 | 1.180397 | 1.111374 | 1.158604 | 1.151919 | 1.143431 |
| rho | 0.308044 | 0.307889 | 0.297518 | 0.272961 | 0.289788 | 0.287412 | 0.284392 |
| p | 0.000000 | 0.000000 | 0.000000 | 0.000000 | 0.000000 | 0.000000 | 0.000000 |
| bic | 2.97e+02 | 2.93e+02 | 2.93e+02 | 2.90e+02 | 2.95e+02 | 2.99e+02 | 3.04e+02 |

\* $p < 0.05$ , \*\* $p < 0.01$ , \*\*\* $p < 0.001$ ; (U) = university level variable; ExIn = “excellence initiative”

**Tab. 9d:**
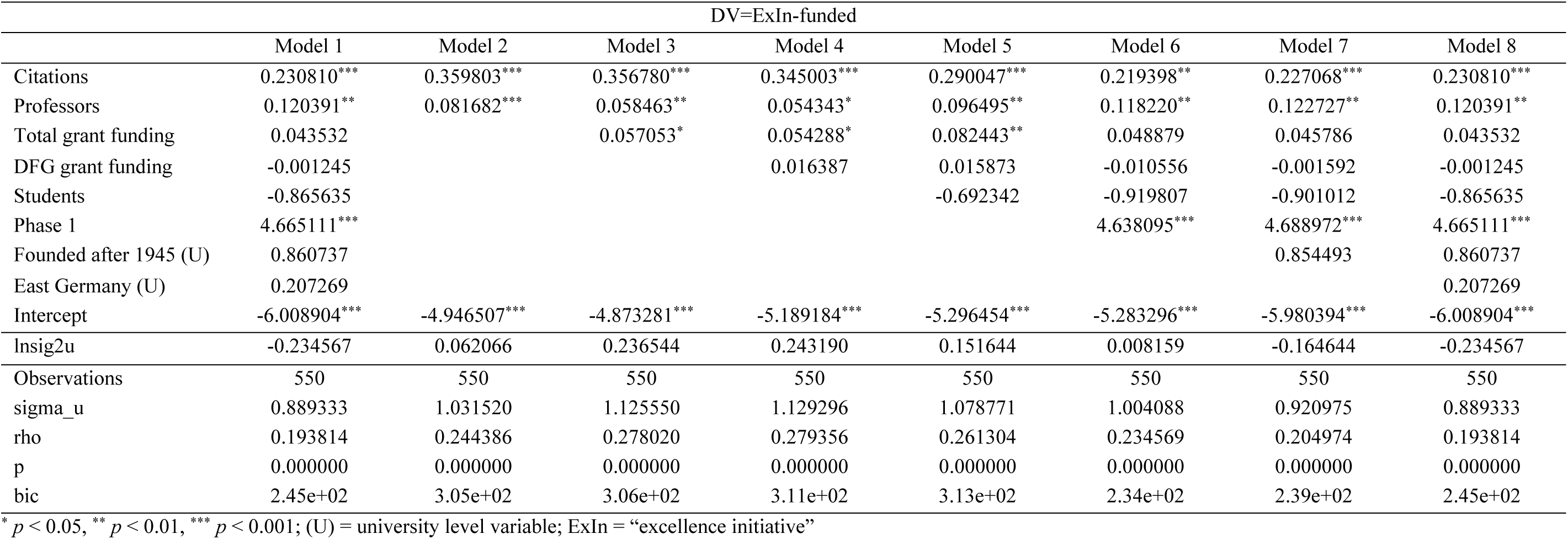
Two-level logistic regression, second “initiative” phase (2012-2017), 12 subject fields with good WoS coverage, all universities.

## Appendix 10: Goodness of fit test

**Tab. 10a:** Goodness of fit test 2006-2011 (Hosmer-Lemeshow)

| Group | Prob | Obs 1 | Exp 1 | Obs 0 | Exp 0 | Total |
| --- | --- | --- | --- | --- | --- | --- |
| 1 | 0.008 | 0 | 1.7 | 239 | 237.3 | 239 |
| 2 | 0.010 | 0 | 2.1 | 239 | 236.9 | 239 |
| 3 | 0.013 | 2 | 2.7 | 237 | 236.3 | 239 |
| 4 | 0.016 | 0 | 3.4 | 239 | 235.6 | 239 |
| 5 | 0.020 | 5 | 4.3 | 233 | 233.7 | 238 |
| 6 | 0.026 | 6 | 5.4 | 233 | 233.6 | 239 |
| 7 | 0.034 | 4 | 7.1 | 235 | 231.9 | 239 |
| 8 | 0.048 | 11 | 9.7 | 228 | 229.3 | 239 |
| 9 | 0.084 | 22 | 14.6 | 217 | 224.4 | 239 |
| 10 | 0.996 | 52 | 51.1 | 186 | 186.9 | 238 |

Number of observations = 2,388
Number of groups = 10
Hosmer-Lemeshow chi2(8) = 13.14
Prob > chi2 = 0.1071

**Tab. 10b:** Goodness of fit test 2012-2017 (Hosmer-Lemeshow)

| Group | Prob | Obs 1 | Exp 1 | Obs 0 | Exp 0 | Total |
| --- | --- | --- | --- | --- | --- | --- |
| 1 | 0.004 | 0 | .6 | 240 | 239.4 | 240 |
| 2 | 0.005 | 0 | 1.1 | 240 | 238.9 | 240 |
| 3 | 0.005 | 1 | 1.2 | 239 | 238.8 | 240 |
| 4 | 0.006 | 1 | 1.3 | 238 | 237.7 | 239 |
| 5 | 0.006 | 0 | 1.5 | 239 | 237.5 | 239 |
| 6 | 0.008 | 1 | 1.7 | 239 | 238.3 | 240 |
| 7 | 0.011 | 2 | 2.2 | 238 | 237.8 | 240 |
| 8 | 0.018 | 3 | 3.3 | 236 | 235.7 | 239 |
| 9 | 0.045 | 9 | 6.7 | 231 | 233.3 | 240 |
| 10 | 1.000 | 108 | 105.6 | 131 | 133.4 | 239 |

Number of observations = 2,396
Number of groups = 10
Hosmer-Lemeshow chi2(8) = 4.49
Prob > chi2 = 0.8100

## Appendix 11: Robustness checks

**Tab. 11a:**
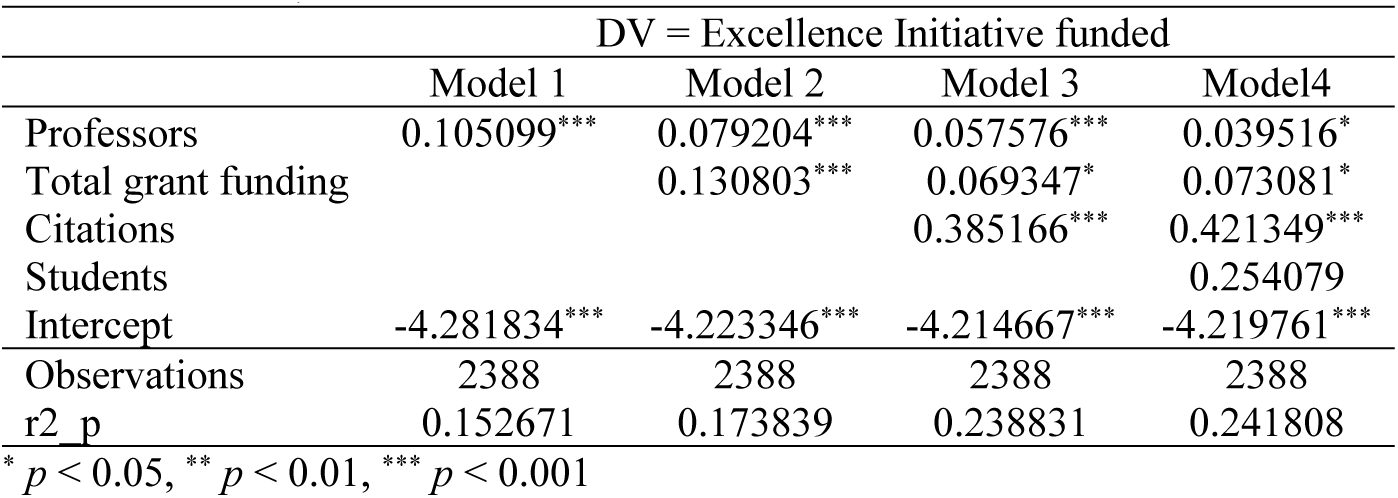
Robustness check. Logistic regression, first “initiative” phase (2006–2011), all universities, incl. citations.

| DV = Excellence Initiative funded |  |  |  |  |
| --- | --- | --- | --- | --- |
|  | Model 1 | Model 2 | Model 3 | Model4 |
| Professors | 0.105099*** | 0.079204*** | 0.057576*** | 0.039516* |
| Total grant funding |  | 0.130803*** | 0.069347* | 0.073081* |
| Citations |  |  | 0.385166*** | 0.421349*** |
| Students |  |  |  | 0.254079 |
| Intercept | -4.281834*** | -4.223346*** | -4.214667*** | -4.219761*** |
| Observations | 2388 | 2388 | 2388 | 2388 |
| r2_p | 0.152671 | 0.173839 | 0.238831 | 0.241808 |

\* $p < 0.05$ , \*\* $p < 0.01$ , \*\*\* $p < 0.001$

**Tab. 11b:** Robustness check. Logistic regression, second “initiative” phase (2012–2017), all universities, incl. citations.

| DV = Excellence Initiative funded |  |  |  |  |  |
| --- | --- | --- | --- | --- | --- |
|  | Model 1 | Model 2 | Model 3 | Model 4 | Model 5 |
| Professors | 0.127277*** | 0.100775*** | 0.077238*** | 0.116906*** | 0.124363*** |
| Total grant funding |  | 0.083858*** | 0.037940 | 0.057827** | 0.014639 |
| Citations |  |  | 0.267022*** | 0.211486*** | 0.201762*** |
| Students |  |  |  | -0.653553** | -0.916477* |
| Phase I |  |  |  |  | 5.867898*** |
| Intercept | -4.410611*** | -4.309952*** | -4.312490*** | -4.348694*** | -5.298393*** |
| Observations | 2396 | 2396 | 2396 | 2396 | 2396 |
| r2_p | 0.216103 | 0.231675 | 0.301258 | 0.310636 | 0.651632 |

\* $p < 0.05$ , \*\* $p < 0.01$ , \*\*\* $p < 0.001$

## Footnotes

1 A peculiarity of the second phase is its extension into the year 2019, before the start of “Excellence Strategy,” the “initiative“’s successor; therefore, all university subjects (and universities via the IS funding line) received additional funding before the “Excellence Strategy” became operational late in 2019.

2 When interpreting Table 3a, note that the “no funding” category includes subject fields in all three university categories mentioned above (Table 2). In other words, there are also subject fields at “excellence” universities that were not funded by the “initiative.”

